# The centriolar satellite protein Cfap53/Ccdc11 facilitates the formation of the first zygotic microtubule organizing center in the zebrafish embryo

**DOI:** 10.1101/2020.11.18.388652

**Authors:** Sven Willekers, Federico Tessadori, Babet van der Vaart, Heiko Henning, Riccardo Stucchi, Maarten Altelaar, Bernard A.J. Roelen, Anna Akhmanova, Jeroen Bakkers

## Abstract

In embryos from most animal species a zygotic centrosome is assembled by the centriole derived from the sperm cell and pericentriolar proteins present in the oocyte. This zygotic centrosome acts as a microtubule organizing center (MTOC) to assemble the mitotic spindle in the first and all subsequent cell divisions. As MTOC formation has been studied mainly in adult cells, very little is known about the formation of the first zygotic MTOC. Here we find that zebrafish (*Danio rerio*) embryos lacking maternal or paternal Cfap53, a centriolar satellite protein, arrest during the first cell cycle due to a failure in proper formation of the mitotic spindle. During the first cell cycle Cfap53 co-localizes with γ-tubulin and other centrosomal and centriolar satellite proteins to the very large MTOC. Furthermore, we find that γ-tubulin localization to the MTOC is impaired in the absence of Cfap53 or when the microtubule network is disrupted. Based on these results we propose a model in which maternal and paternal Cfap53 participates in the organization of the first zygotic MTOC of the embryo. Once the zygotic MTOC is formed, Cfap53 is dispensable for MTOC formation and integrity in subsequent cell divisions.

## Introduction

The animal embryo must undergo multiple rounds of cell divisions to form a multicellular organism composed of hundreds of different cell types. During each cell division microtubule organizing centers (MTOC) participate in the formation of the mitotic spindle required to segregate chromosome into the two daughter cells. In dividing animal cells the centrosome is the main MTOC. Centrosomes are composed of a pair of centrioles surrounded by a complex protein structure consisting of microtubule nucleating and anchoring proteins called the pericentriolar material (PCM) (reviewed in Doxsey, 2001; Gould and Borisy, 1977).

In most animal oocytes centrosomes are eliminated during oogenesis (Schatten et al., 1989; Szollosi et al., 1972). After fertilization, centrosomes are reassembled, which requires the interaction between the centrioles present in the sperm cell and maternal factors present in the oocyte (Holy and Schatten, 1997; Yabe et al., 2007). The sperm cell contains a pair of centrioles termed proximal and distal centrioles, but most centrosomal components are eliminated from the sperm cell (reviewed in Schatten, 1994; Sutovsky and Schatten, 1999). Subsequently, after fertilization the proximal centriole will recruit centrosomal components that are deposited maternally in the oocyte to form a functional zygotic centrosome (reviewed in Schatten, 1994). One of the best characterized centrosomal components that is recruited to the MTOC after fertilization is *γ*-tubulin. It is part of the γ-tubulin ring complex (γTuRC) that serves as a template for nucleating microtubules (reviewed in Kollman et al., 2011). How γTuRC recruitment is facilitated towards the MTOC is still unclear. *In vitro* and cell culture studies have shown that both microtubule-based active transport and passive diffusion contribute to centrosomal γTuRC recruitment (Dammermann and Merdes, 2002; Félix et al., 1994; Khodjakov and Rieder, 1999; Moritz et al., 1998; Quintyne et al., 1999; Schnackenberg et al., 1998; Stearns and Kirschner, 1994; Young et al., 2000). In rodents the sperm centriole is lost during spermatogenesis (Schatten et al., 1986; Woolley and Fawcett, 1973). However, MTOCs are still formed by recruitment of centrosomal components independently of centrioles (Gueth-Hallonet et al., 1993).

Zebrafish embryos during early cleavage stages have large MTOCs in which PCM components do not form a well-defined structure, but appear as cytoplasmic foci (Lindeman and Pelegri, 2012; Rathbun et al., 2020). This specific organization of the MTOC could be a mechanism that enables the mitotic spindle to span the large embryonic cells. The establishment of this unique MTOC after fertilization is probably highly coordinated and dependent on maternal and/or paternal factors present in the fertilized embryo. Forward genetic screens in zebrafish is an unbiased approach to identify parental factors that have an important biological role in early development (Pelegri and Mullins, 2016). Several zebrafish and medaka mutants have been generated via this approach and proven to be very informative for the identification and characterization of some of these maternal and paternal factors (Abrams et al., 2020; Dekens et al., 2003; Inoue et al., 2017; Yabe et al., 2007). The zebrafish gene *cellular atoll (cea)* encodes for the centriole duplication factor Sas-6 (Yabe et al., 2007). Maternal *cea/sas-6* mutant embryos proceed through the first cell division but show mitotic defects from the second cell division onward. Paternal *cea/sas-6* mutant embryos have a delayed first cell division, but cell division proceeds normally thereafter. The zebrafish gene *futile cycle* encodes for Lymphoid-restricted membrane protein (Lrmp) and in maternal *fue/lrmp* mutant embryos the male and female pronuclei fail to fuse (Dekens et al., 2003; Lindeman and Pelegri, 2012). The two pronuclei fuse to one zygote nucleus in a process called ‘nuclear congression’. Nuclear congression requires the formation of a microtubule aster that nucleates from the proximal centriole and attaches to the pronuclei to bring them together, a process that fails in *fue/lrmp* mutants (Dekens et al., 2003; Lindeman and Pelegri, 2012). In mouse embryos, nuclear congression appears different as it is facilitated by two bipolar spindles formed by two clusters of MTOCs around each pronucleus (Reichmann et al., 2018). The medaka (*Oryzias latipes*) maternal WD40 repeat-containing protein, Wdr8, (the orthologue of human WRAP73), is important during MTOC assembly (Inoue et al., 2017). Maternal zygotic *wdr8* mutant embryos form multipolar mitotic spindles resulting in chromosome alignment errors. With its WD40 containing domain, Wdr8 interacts with the centriolar satellite protein SSX2IP (Inoue et al., 2017). Recently, additional maternal-effect mutations have been identified via a forward genetic screen that effect cell division in the early embryo, expanding the molecular-genetic understanding of parental contribution in early embryonic development further (Abrams et al., 2020).

Centriolar satellites have been identified as non-membranous granules that surround the centrosomes (Dammermann and Merdes, 2002; Kubo et al., 1999; Odabasi et al., 2019; Prosser and Pelletier, 2020). The best known marker for centriolar satellites is PCM1 as it is the first centriolar satellite protein that was identified (Balczon et al., 1994; Kubo et al., 1999; Kubo and Tsukita, 2003). Since then many more proteins have been identified as components of centriolar satellites, including the cilia- and flagella- associated protein Cfap53/Ccdc11 (hereafter referred to as Cfap53) (Silva et al., 2016). Many of the identified centriolar satellites contain coiled-coil domains, including PCM1 and Cfap53 (Balczon et al., 1994; Kubo et al., 1999; Silva et al., 2016).

To gain insight into the formation of the MTOC and the role of centriolar satellite proteins herein, we studied the zebrafish *cfap53* mutant. We and others previously showed that the zebrafish *cfap53* gene is expressed in ciliated organs of the embryo, including Kupffer’s vesicle, which is the zebrafish left-right organizer (Bakkers et al., 2009; Hirokawa et al., 2006). *Cfap53* mutant embryos develop normally but show organ laterality defects due to compromised cilia function in the Kupffer’s vesicle, which resembles patients with homozygous *CFAP53* mutations (Narasimhan et al., 2015; Noël et al., 2016; Perles et al., 2012; Silva et al., 2016). Adult homozygous *cfap53* mutants are viable. In this study we analyze their offspring and find that maternal and paternal Cfap53 is important during the first cell cycle. Cfap53 interacts with centriolar satellite proteins and centrosomal components and plays a role in the localization of γ-tubulin to the zygotic MTOC. In the absence of both maternal and paternal Cfap53, the formation of the first zygotic MTOC is impaired in most embryos, which affects spindle formation and causes a cell cycle arrest. Embryos that do form a functional MTOC and spindle in the absence of Cfap53, progress through subsequent cell divisions without apparent cell cycle defects. These findings reveal a novel role for Cfap53 during zygotic MTOC formation in the zebrafish embryo.

## Results

### Cfap53 has a maternal and paternal role in the first embryonic cell division

When homozygous *cfap53−/−* fish were crossed we observed that 67% of the progeny did not develop into normal embryos but instead arrested at the first cell cleavage with a single nucleus (Fig. 1*A,B*). In time-lapse movies of these embryos lacking maternal (M) and paternal (P) *cfap53* (hereafter referred to as MP*cfap53−/−)* we observed temporarily invaginations of the cell membrane, but these never resulted in cell divisions. Arrested MP*cfap53−/−* embryos displayed an aberrant nuclear morphology as staining of the nuclear DNA with DAPI showed both enlarged as well as micronuclei (Fig. 1*D,E*). Importantly, the MP*cfap53−/−* embryos in which the first cell division did occur, developed normally and could be grown to viable and fertile adults (Fig. 1*A*). Interestingly, embryos derived from only homozygous *cfap53−/−* males or females crossed to an otherwise wild type female or male respectively, displayed the same arrest in the first cell division, although their frequency was reduced (Fig. 1*C*). Interestingly, the paternal effect is larger than the maternal effect. As unfertilized oocytes arrest in a similar manner, we investigated if the arrested MP*cfap53* −/− embryos were fertilized. We first analyzed sperm cell number and motility of *cfap53−/−* males but we did not observe any significant differences with sperm isolated from wild type males (Fig. 2*A* & *B*, Table S1). After fertilization the male and female pronuclei come together in a process referred to as ‘nuclear congression’, after which they fuse to form the zygotic nucleus. When analyzing nuclear fusion at 15 minutes post fertilization (mpf) in MP*cfap53−/−* embryos, we found no significant difference with wild type embryos (Fig. 2*C*). These results indicate that nuclear congression is not affected in MP*cfap53−/−* embryos and suggests that most arrested embryos were fertilized. To confirm this, we analyzed whether DNA replication was initiated, as this occurs only in fertilized eggs (Dekens et al., 2003). Corroborating our hypothesis, we found that a large proportion of arrested MP*cfap53−/−* embryos efficiently incorporate EdU (Fig. 2*D* and *E*). From these results we conclude that MP*cfap53−/−* embryos are fertilized, initiate the cell cycle but are arrested in the first cell division. These results suggest a role for Cfap53 in cell cycle progression rather than cell cycle initiation.

**Figure 1.**
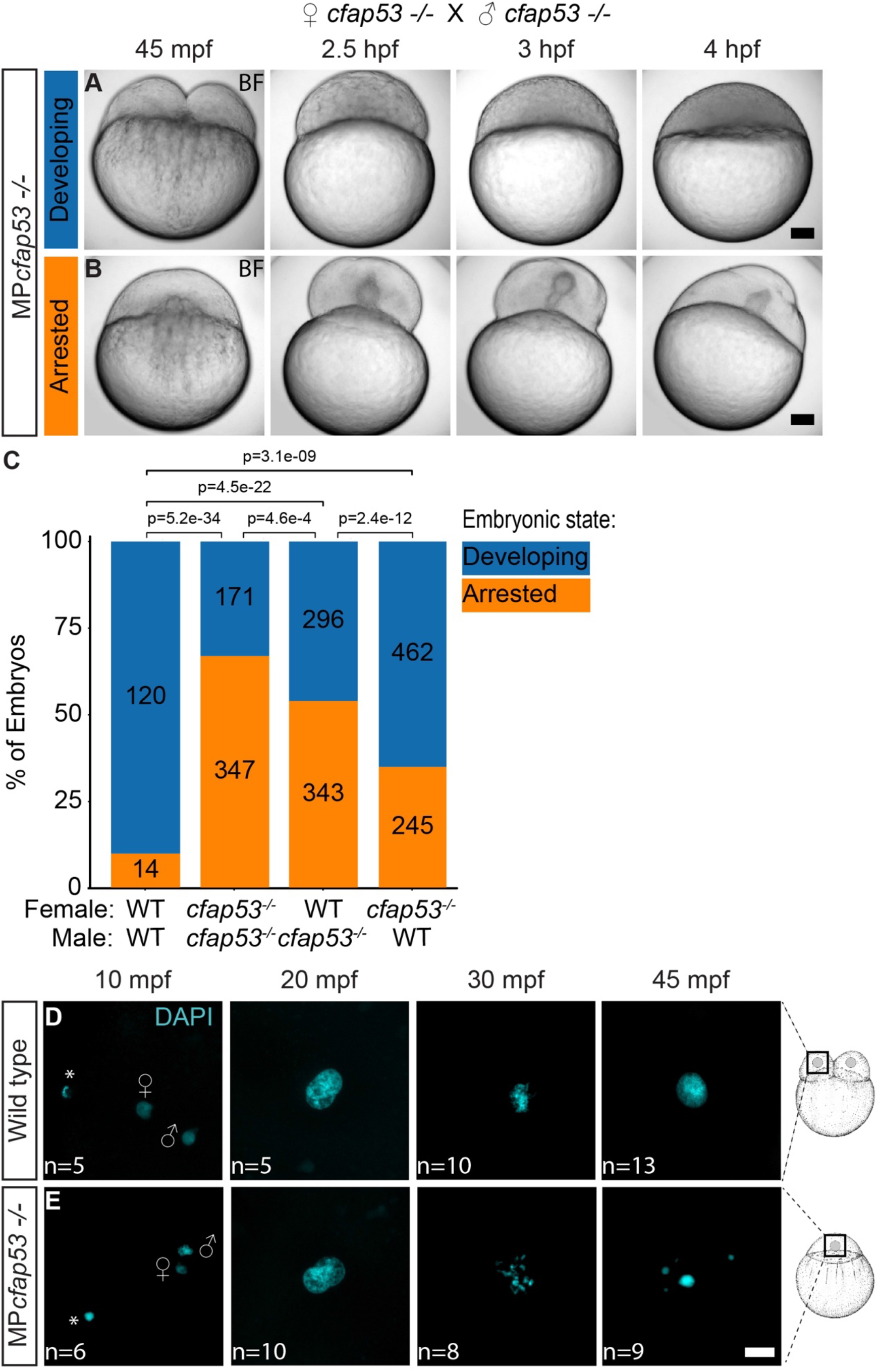
MPcfap53 −/− embryos arrest and display aberrant nuclear morphology. (A,B) Time series of maternal and paternal (MP)*cfap53* −/− embryos from 45 mpf up to 4 hpf. While a small fraction of the MP*cfap53* −/− embryos display normal development (A, n=15), most embryos arrested at the 1-cell stage (B, n=52). (C) Bar plot showing the distribution of arrested (orange) and developing (blue) embryos in clutches of embryos derived from parents with the indicated genotypes. (D,E) Confocal images of wild type (D) and MP*cfap53* −/− embryos (E) at indicated stages of development stained with DAPI to visualize nuclei, indicating aberrant nuclear morphology starts to form at 30 mpf. Asterisk indicates polar bodies. Gender symbols indicate male or female pro-nucleus. Chi-squared test was used to test for significance (p-value <= 0.05). Scale bars indicate 100 microns (A,B) and 20 microns (D,E). Schematics of embryonic development are adapted from Kimmel et al., 1995 (Kimmel et al., 1995).

**Figure 2.**
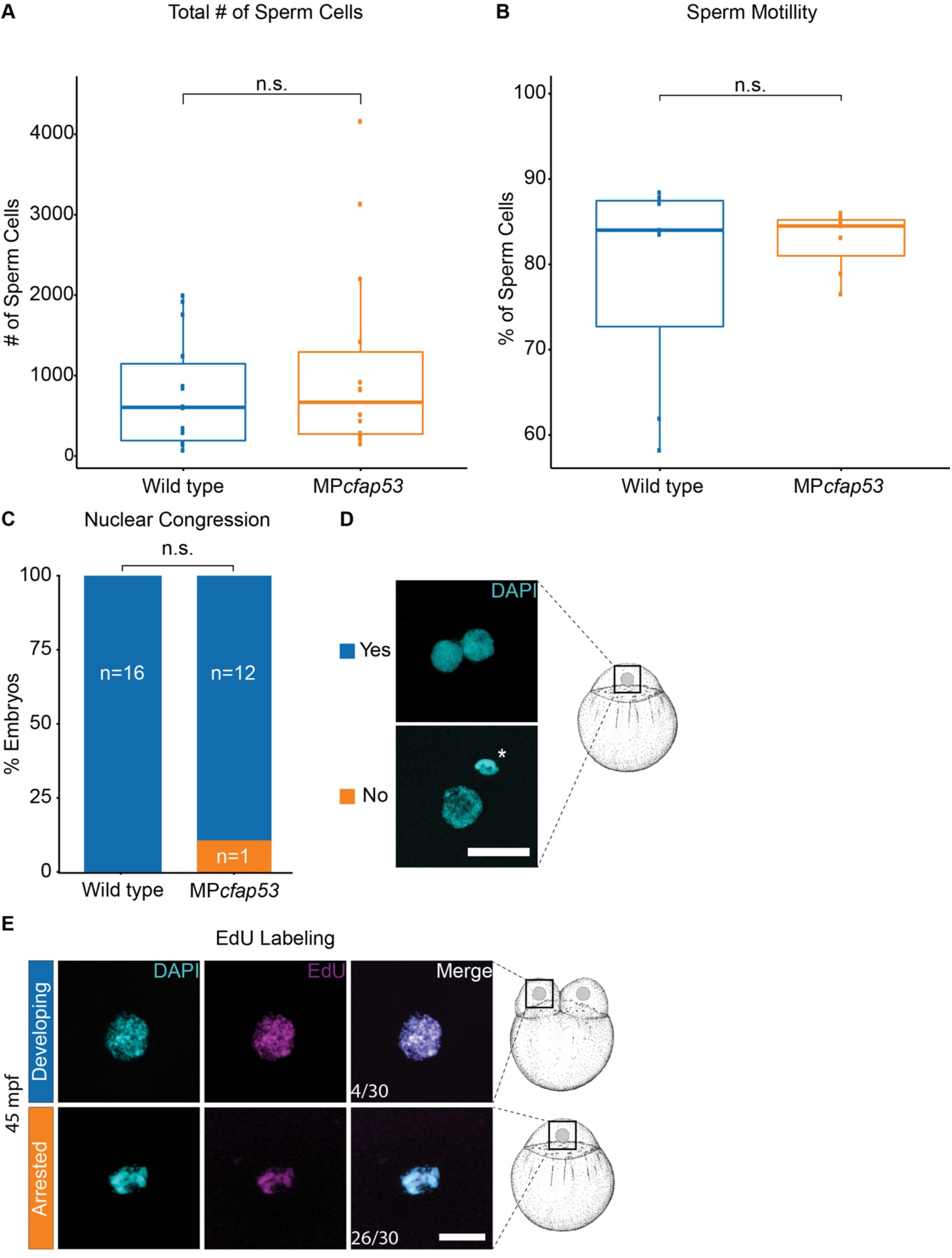
*Cfap53* −/− fish are fertile. (A,B) Boxplots for sperm cell analysis showing the total number of sperm cells counted (A) and percentage of motile sperm cells (B) in spermatozoa derived from wild type (n=7) and *cfap53−/−* males (n=7). (C,D) Quantification of observed nuclear congression in wild type (n=16) and MP*cfap53* −/− (n=13) embryos (C). Confocal images of 1-cell stage embryos stained with DAPI showing either lack of nuclear congression or initiation of nuclear congression of male and female pronuclei (asterisk indicates polar body) (D). (E, n=30) Confocal images showing DAPI staining and EdU incorporation in MP*cfap53* −/− embryos at 45 mpf. Chi-squared test was used to test for significance (p-value <= 0.05). Scale bars indicate 20 microns. Schematics of embryonic development are adapted from Kimmel et al., 1995 (Kimmel et al., 1995).

### Cfap53 interactome

Coiled-coil domain containing proteins often act as scaffolds for large protein complexes, which is consistent with the observation that in cultured cells Cfap53 protein localizes to centriolar satellites and is required for their integrity (Silva et al., 2016). To identify proteins that interact with Cfap53 we performed Cfap53 protein pull downs followed by mass spectrometry (MS). As zebrafish embryos contain relatively large amounts of yolk proteins compared to cytoplasmic proteins that could hamper such an approach, we instead expressed two fusion proteins containing N-terminal or C-terminal biotinylated GFP (bioGFP) and human CFAP53 in HeLa and HEK293 cells. In HeLa cells, zebrafish BioGFP-Cfap53 localized to centrosomes, marked by γ-tubulin, and cytoplasmic aggregates around the centrosomes, reminiscent of centriolar satellites (Fig. S1). Streptavidin-based pull downs of both fusion proteins co-expressed with biotin ligase BirA or biotin ligase BirA expression alone as a negative control followed by MS resulted in the retrieval of in total 1148 overlapping proteins. For these two biological samples, the Pearson correlation of the MS scores across all proteins that were detected in both pull downs was 0.71, demonstrating high reliability of the data. Next, the MS scores from both experiments were combined and used for further statistical analysis using the Significance Analysis of INTeractome (SAINT) and fold change calculation (Choi et al., 2011; Mellacheruvu et al., 2013) (see detailed description in Materials and Methods). From this extensive analysis 88 proteins were considered putative binding partners of Cfap53 (Table S2). Network and GO analysis enabled us to group these proteins into functional classes (Fig. 3*A*, S2).

**Figure 3.**
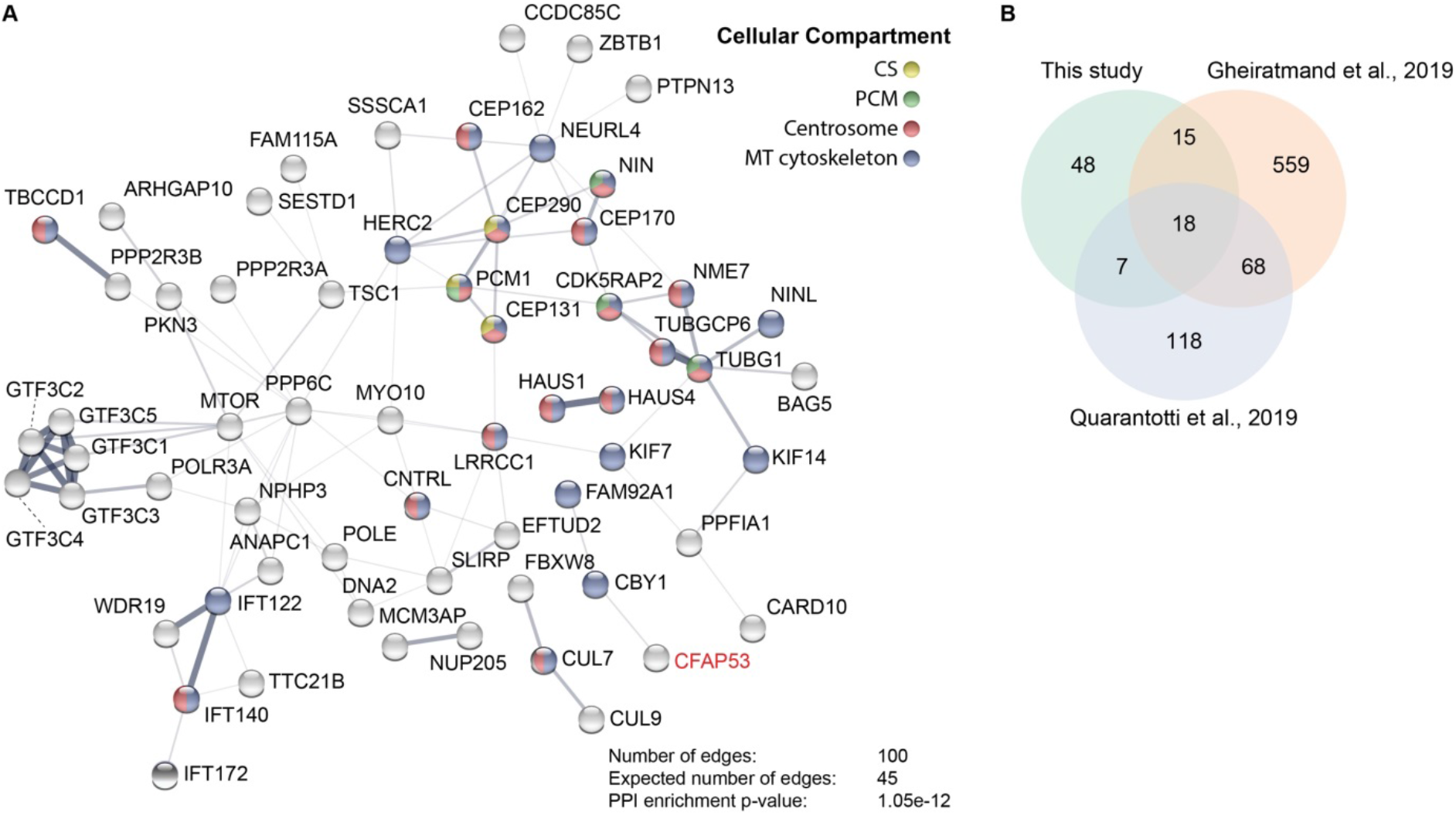
Cfap53 interacts with centrosomal, centriolar satellite and microtubule associated proteins. (A) Network analysis using String-DB (Szklarczyk et al., 2015) from proteins identified by MS analysis after Streptavidin-based purification using bioGFP-Cfap53 as a bait. Only high confident hits from the MS dataset were used which were obtained via stringent filtering steps on raw data (see Materials and Methods). Proteins shows are proteins with a significantly higher number of edges (representing protein-protein associations) than what is expected stochastically. The thickness of the lines between the nodes do not represent strength or specificity of a given interaction, but represents the approximate confidence on a scale from zero to one of a found association being true, given the available evidence. Node colors represent cellular compartment of the indicated protein. (B) Venn diagram showing number of proteins identified in this study by bioGFP-Cfap53 pulldown with MS analysis overlapping with the centriolar satellite proteins that were identified by Gheiratmand et al. and Quarantotti et al. (Gheiratmand et al., 2019; Quarantotti et al., 2019).

A large number of proteins are known for their localization in centrosomes and centriolar satellites, such as PCM1, NIN, CEP131, CEP290 and CEP170, which is consistent with the observed colocalization of Cfap53 and PCM1 (Balczon et al., 1994; Kim et al., 2008; Kubo et al., 1999; Kubo and Tsukita, 2003; Quarantotti et al., 2019; Silva et al., 2016; Staples et al., 2012; Stowe et al., 2012). PCM1 acts as a scaffold for centriolar satellite assembly and facilitates centrosomal protein trafficking (Balczon et al., 1994; Kubo et al., 1999; Kubo and Tsukita, 2003). We therefore compared our protein list with a published list of 211 satellite proteins resulting from an affinity purification of satellites using PCM1 as a bait (Gheiratmand et al., 2019; Quarantotti et al., 2019). We indeed observed a significant overlap with these interactomes as 25 out of 88 proteins (28%, p-value: 1.8e-22) that were enriched in the bioGFP-Cfap53 pull down were also present in the PCM1 affinity purification (Fig. 3*B*). In addition, a proximity-dependent biotin identification approach with 22 satellite proteins, including Cfap53, identified an interactome of 660 proteins (Gheiratmand et al., 2019). Comparison with the Cfap53 interactome revealed that 33 out of 88 proteins (37%, p-value: 6.4e-18) overlapped with the centrosomal satellite proteome (Fig. 3*B*).

Another prominent group of proteins copurifying with Cfap53 are γTuRC components γ-tubulin (TUBG1) and GPC6, as well as proteins that interact with γTuRC such as, CDK5RAP2 and the kinase NME7 (Choi et al., 2010; Liu et al., 2014; Murphy et al., 2001).

Moreover, the MS data contained components of HAUS complex (HAUS1 and HAUS4), which localizes to spindles and centrosomes and controls branching microtubule nucleation (Goshima et al., 2008; Lawo et al., 2009).

Together these data show that Cfap53 can interact with centrosomal and centriolar satellite proteins as well as proteins with known functions in microtubule nucleation.

### Cfap53 localizes to MTOC during embryonic development

As Cfap53 localization in zebrafish embryos had not been determined we generated a Tg(*ubi:GFP-Cfap53*) line, in which a GFP-Cfap53 fusion protein is expressed ubiquitously including developing oocytes (Fig. 4*A*). To test if the GFP-Cfap53 protein is functional, the Tg(*ubi:GFP-Cfap53*) was crossed into the *cfap53^hu1047^* line. We first incrossed Tg(*ubi:GFP-Cfap53*)/*cfap53 +/-* fish and analysed their progeny for laterality defects, as we previously described laterality defects in *cfap53*−/− embryos (Noël et al., 2016) (Fig. S3). We found that the laterality defect was rescued in *cfap53−/−* embryos expressing the GFP-Cfap53 fusion protein. Next we crossed adult *cfap53−/−* fish that carried the Tg(*ubi:GFP-Cfap53*) and their progeny was scored for GFP-Cfap53 expression and embryonic development. Importantly, the GFP-Cfap53 was also able to rescue the maternal *cfap53*−/− phenotype as the percentage of arrested embryos in MP*cfap53*−/− embryos dropped from 67% to 20% when maternal GFP-Cfap53 was present (Fig. S3). From these results we concluded that GFP-Cfap53 is a functional protein and therefore could be used to determine its subcellular localization.

**Figure 4.**
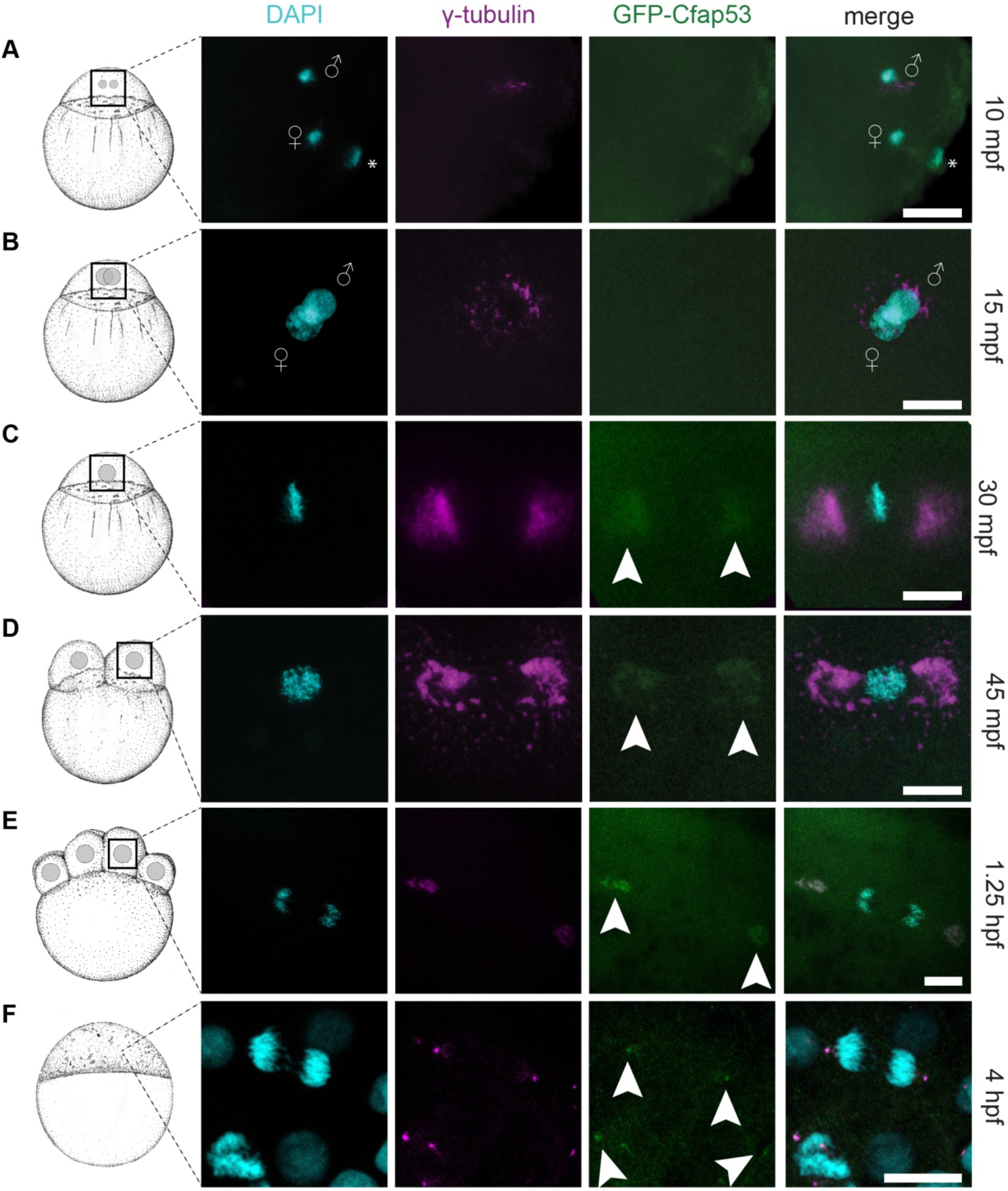
GFP-Cfap53 localization during cleavage stages. (A-F) Maximal projections of confocal stacks from Tg(*ubi:GFP-cfap53*) embryos immunolabelled for GFP, γ-tubulin and DAPI from 30 mpf until 4 hpf. At 10 mpf male and female pronuclei and female polar body are visible. Some γ-tubulin foci around male pronucleus can been observed, while GFP-Cfap53 has a diffuse localization (A, n=7). At 15 mpf both pronuclei are almost fused and some γ-tubulin can be observed around the nuclei, while GFP-Cfap53 localization is still diffuse (B, n=7). At 30 mpf chromosomes have aligned along the metaphase plate and strong γ-tubulin staining can be observed on either side of it. At this moment, also GFP-Cfap53 starts to localize to both sides of the metaphase plate where it colocalizes with γ-tubulin (C, n=12). At 45 mpf the first cell division has completed and one of the two nuclei is shown. Both GFP-Cfap53 and γ-tubulin are visible in broad domains on both sides of the nucleus during prophase (D, n=8). Single cell of a 8-cell stage blastula with segregating chromosomes during telophase of mitosis. GFP-Cfap53 and γ-tubulin co-localize in a more condensed region at both sites of the division plane (E, n=7). Blastula cell in telophase during mitosis at the sphere stage (4 hpf). GFP-Cfap53 and γ-tubulin co-localize to a confined region on both sides of the division plane (F, n=1). Asterisk indicates polar body. Gender symbols indicate male or female pro-nucleus. Arrowheads indicate localized GFP signal colocalizing with γ-tubulin. Scale bars indicate 20 microns. Schematics of embryonic development are adapted from Kimmel et al., 1995 (Kimmel et al., 1995).

We investigated the subcellular localization of GFP-Cfap53 in embryos starting at 10 mpf up to 4 hpf and compared it to γ-tubulin localization as a marker for the MTOC (Lindeman and Pelegri, 2012; Rathbun et al., 2020; Yabe et al., 2007). During the first 15 minutes of development we observed a weak GFP-Cfap53 signal equally distributed throughout the cytoplasm while some γ-tubulin was localized around the male pronucleus (Fig.4*A* & *B*). At 30 mpf, when the first mitosis of the zygote has started, we observed accumulation of GFP-Cfap53 on either side of the metaphase plate containing the aligned chromosomes. At this stage we observed for the first time a colocalization of GFP-Cfap53 with γ-tubulin (Fig. 4*C*). As embryo development proceeds with a new cell division every 15 minutes, both GFP-Cfap53 and γ-tubulin localization became more confined into two structures on either side of the aligned or segregating chromosomes (Fig. 4*E-F*). Together these results indicate that in the oocyte GFP-Cfap53 is initially diffusely localized throughout the cytoplasm and that after pronuclear fusion both Cfap53 and γ-tubulin accumulate to form large MTOCs consisting of multiple protein foci. As we observed that paternal Cfap53 is required to initiate the first cell division, we investigated GFP-Cfap53 localization in sperm cells. We observed that the GFP-Cfap53 localizes near the sperm DNA in two domains. The first domain is around the centrioles marked by Centrin and the second domain is in a single spot of unknown origin near the centrioles (Fig. S4).

### Cfap53 facilitates the formation of the MTOC after fertilization

To confirm that GFP-Cfap53 localizes to the MTOC in the zebrafish embryo, we co-stained GFP-Cfap53 expressing embryos with antibodies that recognizes Centrin and γ-tubulin, two key components of MTOCs. Indeed, we observed colocalization of GFP-Cfap53 with both Centrin and γ-tubulin (Fig. 5*A-C*).

**Figure 5.**
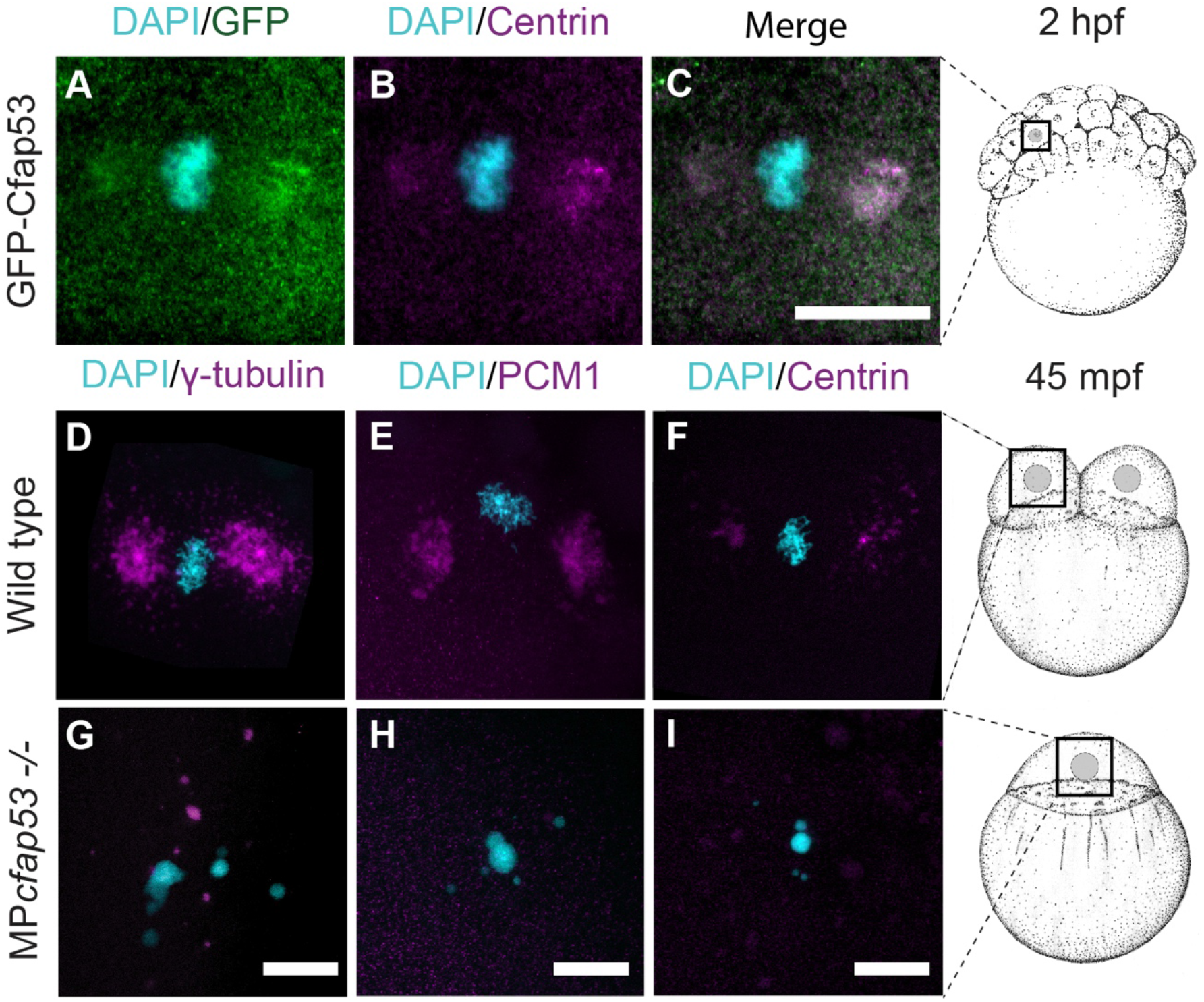
Cfap53 is important for the localization of centrosomal components and centriolar satellites at the MTOC. (A-C, n=9) Maximal projections of confocal stacks from fixed Tg(ubi:*GFP-cfap53)* transgenic embryos immunolabeled for Centrin at 2 hpf shows strong colocalization of GFP-Cfap53 with Centrin during metaphase. (D-I) Maximal projections of confocal stacks from fixed wild type or MP*cfap53 −/−* embyos at 45mpf immunolabeled for centrosomal components. γ-tubulin (D, n=8, G, n=15), PCM1 (E, n=7, H, n=15), and Centrin (F, n=5, I, n=8) in wildtype embryos (D-F) or MP*cfap53* −/− non-developing embryos (G-I) at 45 mpf fixed during metaphase. γ-tubulin, PCM1 and Centrin accumulate at both sides of the division plane in wild type embryos. In MPcfap53 −/− embryos γ-tubulin forms large foci and the chromosomes are not alinged. PCM1 and Centrin have a diffuse localization in the MP*cfap53* −/− embryos. Scale bars indicate 20 microns. Schematics of embryonic development are adapted from Kimmel et al., 1995 (Kimmel et al., 1995).

Based on the Cfap53 pull down results and its localization, we examined a possible role of Cfap53 in the formation of the MTOC. As the MS data revealed a possible interaction between Cfap53 with PCM1 and γ-tubulin, we investigated the localization of these and other centrosomal proteins in wild type and MP*cfap53−/−* embryos. In wild type embryos, γ-tubulin, PCM1 and Centrin were localized in the large MTOC structures typical for early cleavage stage zebrafish embryos (Fig 5A-C) (Dekens et al., 2003; Lindeman and Pelegri, 2012; Rathbun et al., 2020). In arrested MP*cfap53−/−* embryos, γ-tubulin signal was present, albeit at reduced levels, and it formed granule-like foci dispersed throughout the cytoplasm (Fig. 5*D*). In addition, PCM1 and Centrin signals were diffuse and no longer accumulated in MTOC structures in arrested MP*cfap53−/−* embryos (Fig. 5*E* & *F*).

Failure in MTOC formation will result in a disorganized microtubule network and cell cycle arrest, since proper bipolar spindles cannot be formed. To test whether the observed arrest in the first cell cycle of MP*cfap53−/−* embryos could be due to defective bipolar spindle formation, we used the Tg(*XlEef1a1:dclk2a-GFP)*, which labels microtubules *in vivo*. As the Dclk2a-GFP fusion protein is maternally provided we used it to visualize the microtubule network during the first cell division. In developing MP*cfap53* embryos we observed microtubule bundles nucleating from the MTOC to form astral microtubules and a symmetric bipolar spindle (Fig.6*A*). In MP*cfap53−/−* arrested embryos however, the bipolar spindle was not formed, instead we observed that microtubule bundles still nucleate but in a disorganized fashion originating from the chromosomes (Fig. 6*B*), which explains their cell cycle arrest. Taken together, these results indicate that Cfap53 facilitates MTOC formation and therefore the assembly of the mitotic spindle during the first cell division. However, Cfap53 is dispensable for MTOC formation and assembly of the mitotic spindle during subsequent cell divisions.

**Figure 6.**
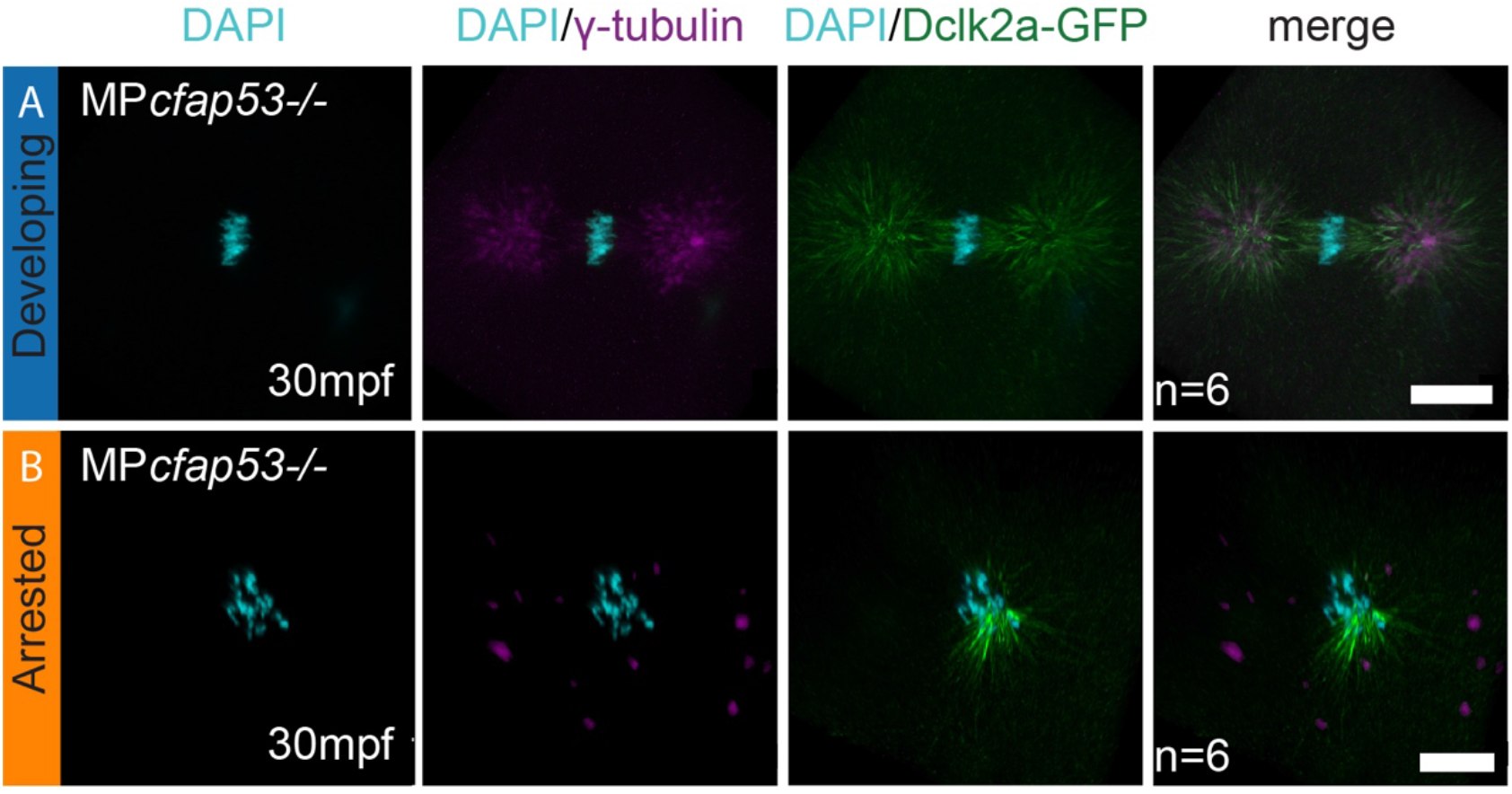
Cfap53 facilitates assembly of the first zygotic spindle. (A,B) Maximal projections of confocal stacks on Tg*(XlEef1a1:dclk2a-GFP)*/MP*cfap53* −/− embryos at 30 mpf and immunolabeled for GFP, γ-tubulin and DAPI. While six embryos formed centrosomes and a normal mitotic spindle during the first zygotic cell division (A, n=6), six embryos failed to assemble the centrosomes and the mitotic spindle (B, n=6). Instead, microtubule originating from the chromosomes is observed in these embryos. Scale bars indicate 20 microns.

### The microtubule network is important for the first zygotic MTOC formation

When analyzing the Dlck2a-GFP and γ-tubulin colocalization more closely, we noticed that γ-tubulin forms foci varying in size at apparent microtubule ends or dispersed along microtubules. The Dclk2a-GFP and γ-tubulin did not appear to colocalize (Fig. 7*A* & *B*), which was confirmed by a very low Pearson’s coefficient of 0.0997 for Dclk2a-GFP and γ-tubulin colocalization (Fig. 7*B*). This observation suggests that during the first cell division γ-tubulin is associated with the microtubules similar to what has been shown for PCM1 (Kubo and Tsukita, 2003) Therefore, we investigated whether microtubules are required for γ-tubulin localization at the MTOC during the first cell division. We therefore treated wild type embryos with nocodazole or taxol to either depolymerize or stabilize microtubules respectively, in developing embryos (Fig. 7C). While taxol treatment arrested the embryos at the metaphase stage during the first mitosis, there was no effect on nuclear morphology or γ-tubulin localization (Fig. *7D*). Interestingly, embryos treated with nocodazole showed aberrant nuclear morphology and dispersed cytoplasmic granules of γ-tubulin, similar to those seen in MP*cfap53−/−* embryos (Fig. 5D and 6*B*). Next, we addressed whether microtubule dependent localization of γ-tubulin also occurs in later cell divisions by treating embryos with nocodazole during the third cell division. Importantly, nocodazole treatment of embryos during the third cell division, resulted in a cell cycle arrest at the 4-cell stage but had no apparent effect on γ-tubulin localization to the MTOC (Fig. 7E-F). To test if the microtubule network is solely involved in this effect, we depolymerized the actin cytoskeleton via chemical treatment of wild type embryos with Latrunculin A. Overall, cell morphology was aberrant, however there was no effect on nuclear morphology or γ-tubulin localization at the MTOC (Fig. 7*C-D*). We therefore conclude that a functional microtubule network is required for the localization of γ-tubulin at the MTOC during the first cell division in the zebrafish embryo.

**Figure 7.**
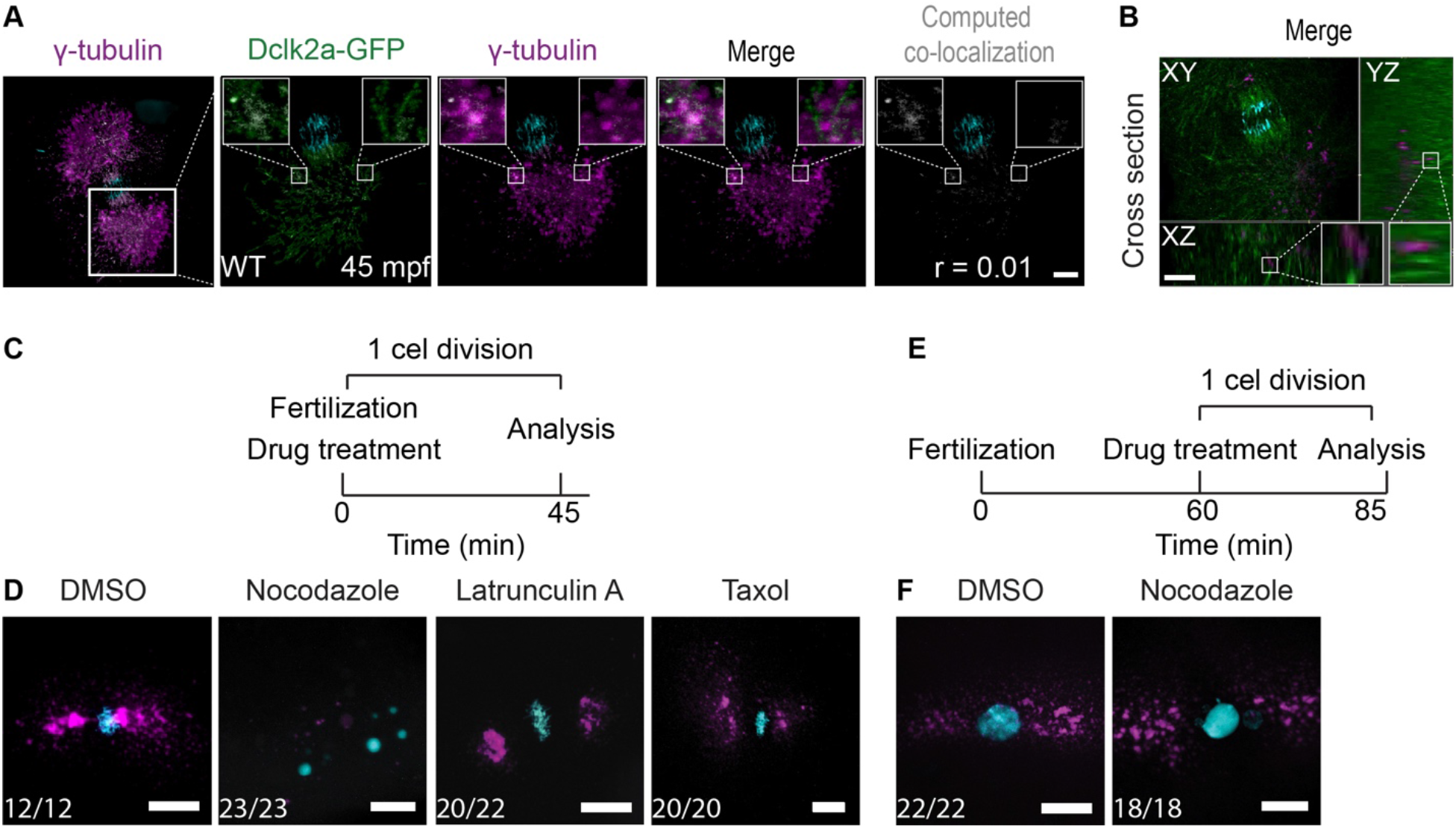
Microtubule-depenend γ-tubulin localization during the first cell division. (A,B) Maximal projections of confocal stacks from Tg*(Xla.Eef1a1:Dclk2a-GFP)* embryo immunolabeled for DAPI (in cyan), Dclk2a-GFP (in green) and γ-tubulin (in magenta) at 45 mpf fixed during telophase. Colocalization of γ-tubulin and Dclk2a-GFP is shown in white in the merge. Computation of colocalization gives an average Pearson’s coefficient of 0.1 indicating that there is very little colocalization of γ-tubulin and Dclk2a-GFP (A, n=5). Optical cross section through 45 mpf embryo immunostained for γ-tubulin and Dclk2a-GFP at the region of one of the centrosomes, which shows that γ-tubulin is localized directly adjacent to the microtubule bundles (B, n=5). Scalebar indicates 100 microns. White squares represent zoom ins of outlined area. Schematic overview of the drug treatment experiment (C), for which the results are shown in (D). Embryos were treated during the first cell division. Maximal projections of confocal stacks of embryos at 45 mpf immonostained with γ-tubulin (in magenta) and DAPI (in cyan) fixed during telophase (D). Nocodazole treatment (n=23) during the first cell division prevents γ-tubulin accumulation at the MTOC. Lantrunculin A (n=20) and taxol (n=20) treatments have no effect on γ-tubulin localization to the MTOC. Schematic overview of the drug treatment experiment (E), for which the results are shown in (F). Embryos were treated during the third cell division and fixed after 15 minutes of treatment during interphase. Maximal projections of confocal stacks of embryos at 85 mpf immonostained with γ-tubulin and DAPI (F). Nocodazole treatment had no effect on γ-tubulin localization at the MTOC (n=18). Scale bars indicate 20 microns.

## Discussion

The first cell division in embryogenesis is different compared to following cell divisions, as it involves formation of the first MTOC from maternal and paternal components. While this is a critical step to assure formation of the first mitotic spindle, which facilitates proper cell division and further development of the embryo, very little is known about its mechanisms and the factors that are involved. In this study, we describe that the centriolar satellite protein Cfap53 plays an important role in the formation of the first zygotic MTOC.

The cell cycle arrest at the first cell division that we observed in maternal and paternal *cfap53−/−* embryos has an incomplete penetrance. The EdU labeling experiment demonstrates that around 87% of fertilized MP*cfap53*−/− embryos go through S-phase but arrest before completing mitosis. The remaining 13% of the embryos divide and continue to develop normally only displaying the zygotic laterality defects described earlier (Narasimhan et al., 2015; Noël et al., 2016; Perles et al., 2012; Silva et al., 2016). This suggests that the loss of Cfap53 during the first cell division can be compensated by other proteins with a similar function. Recent genetic work in zebrafish has shown that mutations that introduce a premature stop codon in the mRNA, which results in mRNA degradation, can trigger a transcriptional adaptation mechanism (Rossi et al., 2015; Sztal et al., 2018; Zhu et al., 2017). This transcriptional adaptation mechanism causes an upregulation of the expression of related genes and thereby (partially) rescues the phenotype. As the *cfap53^hu10478^* allele is a 7 bp deletion, it results in a frameshift and subsequent introduction of a premature stop codon, which may result in mRNA degradation and activation of the transcriptional adaptation mechanism. mRNA sequencing of oocytes from *cfap53* mutant fish could help to identify compensating genes.

Our genetic analysis indicates that both maternal and paternal Cfap53 are important for progression through the first cell division (Fig. 1). Very few genes have been described with both maternal and paternal phenotypes. Another example of a maternal and paternal mutant is the *cao/sas-6* mutant, which results in either arrested or delayed cell divisions depending on the genotype of the male and female (Yabe et al., 2007). Our finding that paternal Cfap53 is important is consistent with the detection of Cfap53 protein in bovine sperm by mass spectrometry (Firat-Karalar et al., 2014) and our own results demonstrating the presence of GFP-Cfap53 protein in zebrafish sperm by microscopy (Fig. S4). Thus, while the sperm cell loses most of its PCM and centriolar satellite proteins, Cfap53 is maintained during spermatogenesis.

After the sperm aster is formed and nuclear congression is completed, GFP-Cfap53 localizes to either side of the fused maternal and paternal pronuclei. This coincides with a clear γ-tubulin accumulation on either side of the nucleus. We and others have observed that PCM1, centrin and γ-tubulin form cytoplasmic accumulations that appear as a large clouds on either side of the nucleus during the first cell divisions of the zebrafish embryo (Dekens et al., 2003; Lindeman and Pelegri, 2012; Rathbun et al., 2020; Stowe et al., 2012). Why these structures that function as MTOCs are so large is not clear, but might have to do with the large size of the embryo (see also below). PCM1-, centrin- and γ-tubulin-containing structures depend on Cfap53 as these were not formed in MP*cfap53* embryos. The biochemical function of Cfap53 is currently unclear. It has three predicted coiled-coil domains, which are low complexity structures that facilitate protein-protein interactions and are found in numerous centriolar satellite proteins (Gheiratmand et al., 2019; Mason and Arndt, 2004; Quarantotti et al., 2019). It is possible that similar to the *C. elegans* centrosome component, the coiled-coil proteins SPD-5, CFAP53 might facilitate MTOC formation by a passive process called liquid-liquid phase separation (Woodruff, 2018; Woodruff et al., 2017).

What could be the mechanism by which Cfap53 recruits γ-tubulin and other components towards the MTOC after fertilization? In general protein targeting to the MTOC is regulated by two different pathways; (1) active microtubule based transport using centriolar satellites and (2) concentration of proteins to the MTOC via passive diffusion (Conkar et al., 2019; Gillingham and Munro, 2000; Hori and Toda, 2017; Khodjakov and Rieder, 1999; Young et al., 2000; Zimmerman and Doxsey, 2000). For a subset of centrosomal proteins passive diffusion is sufficient in somatic cells, which includes γ-tubulin (Khodjakov and Rieder, 1999). However, passive diffusion of γ-tubulin alone may not be sufficient as zebrafish embryos have much larger volumes during the first cell divisions compared to average somatic cells (1.000x in case of the zebrafish). This can be overcome by active directional microtubule- and motor-dependent transport. Several studies have shown that indeed recruitment of γ-tubulin towards the centrosome in somatic cells is facilitated by active transport dependent on the microtubule network (Quintyne et al., 1999; Schnackenberg et al., 1998; Stearns and Kirschner, 1994; Young et al., 2000). Our observation that γ-tubulin-positive clusters align along microtubules (Fig.7*A* and *B*) and our finding that nocodazole treatment of wild type zebrafish embryos directly after fertilization inhibits MTOC localization of γ-tubulin (Fig. 7*C*), is consistent with microtubule dependency of γ-tubulin recruitment. However, nocodazole treatment of embryos that completed the first cell division did not affect γ-tubulin localization to the MTOC in subsequent divisions (Fig. 7*D*), suggesting that active transport is important for the localization of γ-tubulin at the MTOC during the first cell division, but is dispensable at subsequent cell divisions.

How is Cfap53 facilitating the active localization of γ-tubulin at the MTOC? Based on its localization in somatic cells, CFAP53 was classified as a centriolar satellite protein (Silva et al., 2016). Both our MS analysis in somatic cells and the widespread granular localization of GFP-Cfap53 in 2-cell stage zebrafish embryos, which resembles localization of the satellite marker PCM1, is consistent with localization of Cfap53 in centriolar satellites (Fig. 3 & 4*D*, Table S2). Centriolar satellites are membraneless granules containing proteins that associate and move along microtubules in a dynein-dependent manner to deliver proteins to the centrosome (Dammermann and Merdes, 2002; Kubo et al., 1999). We find that γ-tubulin is highly represented in our GFP-Cfap53 MS analysis and we observed the simultaneous accumulation and strong colocalization of GFP-Cfap53 with γ-tubulin at the 2-cell stage (Fig. 3*A* and Fig 4*D*, Table S2), which would suggest a direct role of Cfap53 in γ-tubulin localization to the MTOCs. Alternatively, Cfap53 has a more indirect role in facilitating γ-tubulin localization to the MTOCs. We found NME7 and CDK5RAP2 in our MS data which are both part of the γ-tubulin ring complex and are recruited in a dynein-dependent manner (Fig. 3*A*, Table S2) (Choi et al., 2010; Hutchins et al., 2010; Jia et al., 2013; Liu et al., 2014). Defects in centriolar satellites from MP*cfap53* mutant embryos may therefore result in the absence of other proteins that are required to recruit γ-tubulin. Importantly, the loss of centriolar satellites in somatic cells, by PCM1 depletion, has little effect on the integrity of the centrosome (Gheiratmand et al., 2019), indicating a clear difference in the role of centriolar satellites in fertilized oocytes and somatic cells. This is again consistent with our observation that Cfap53 facilitates the formation of the first zygotic MTOC and is dispensable for MTOC integrity during subsequent cell divisions.

In conclusion, we discovered a novel function for Cfap53 in the formation of the first zygotic MTOC in zebrafish embryos. Whether this function of Cfap53 is conserved in mammalian embryos remains to be investigated.

## Materials and methods

### Zebrafish genetics and strains

The following fish lines were used in this study: Tupfel Longfin (TL), *cfap53^hu10478^* (Noël et al., 2016), Tg(*h2afz:GFP*) (Pauls et al., 2001), Tg(*Xla.Eef1a1:dclk2a-GFP*) (Tran et al., 2012) and Tg(*ubi:GFP-cfap53*). Fish were maintained at 27.5 °C in a 14/10 h light/dark cycle, according to standard laboratory conditions (Aleström et al., 2020). Embryos were collected and staged according to Kimmel *et al*., 1995 (Kimmel et al., 1995). Animal experiments were approved by the Animal Experimentation Committee (DEC) of the Royal Netherlands Academy of Arts and Sciences (KNAW).

### Plasmid Construction and Transgenesis

The following plasmids were generated in this study: bioEGFP-Cfap53, Cfap53-EGFPbio and GFP-Cfap53 containing tol2 sites. *Cfap53* coding sequence (CDS) was obtained by performing PCR on a PCS2 construct containing human *cfap53* cDNA and cloned into the pbioEGFP-C1 and pbioEGFP-N1 constructs using the Gibson cloning system (Noël et al., 2016), which are modified pEGFP-C1 and pEGFP-N1, respectively, in which a linker encoding the sequence MASGLNDIFEAQKIEWHEGGG, which is the substrate of biotin ligase BirA, was inserted in front of or behind the eGFP-encoding sequence.

The zebrafish GFP-Cfap53 construct was obtained from Narasimhan *et al.*, 2014 (Narasimhan et al., 2015) and cloned into the pME-MCS followed by recombination with p5E’-Ubi (Mosimann et al., 2011), p3E-polyA, and pDestTol2pA7 using the multisite gateway cloning strategy (Kwan et al., 2007). The plasmid DNA was injected at 300 ng/μl in the presence of 25 ng Tol2 mRNA for genomic integration and to generate Tg(*ubi:GFP-Cfap53*). At three dpf, healthy embryos displaying robust GFP fluorescence in the cilia were selected and grown to adulthood. Subsequently, founder fish were identified by outcrossing and their progeny grown to adulthood to establish the transgenic line.

### Cell culture

Human embryonic kidney 239T (HEK293T) and HeLA (Kyoto) cell lines were cultured in medium that consisted of 45% DMEM, 45% Ham’s F10, and 10% fetal calf serum supplemented with penicillin and streptomycin. The cell lines were routinely checked for mycoplasma contamination using LT07-518 Mycoalert assay (Lonza). For the immunolabeling experiment HeLa cells were transfected with plasmids using FuGENE 6 (Promega). For streptavidin pull-down assays from HEK293T cells, plasmids were transfected using polyethylenimine (Polysciences).

### Streptavidin pull-down assays

Streptavidin pull-down assays were performed from HEK293T cell lysates by co-expressing biotin ligase BirA with GFP-tagged constructs containing a biotinylation site (bioGFP-Cfap53 and Cfap53-GFPbio) and only expressing biotin ligase BirA as a negative control. Constructs were transfected together into HEK293 cells using polyethylenimine (PEI, Polysciences) with a 24-h incubation time for protein expression. M-280 Streptavidin Dynabeads (Invitrogen) were blocked in a buffer containing 20 mM Tris, pH 7.5, 20% glycerol, 150 mM NaCl, and 10 μg chicken egg albumin followed by three washes with wash buffer containing 20 mM Tris, pH 7.5, 150 mM NaCl, and 0.1% Triton X-100. HEK293T cells were collected in ice-cold PBS followed by lysis on ice in a buffer containing 20 mM Tris, pH 7.5, 150 mM NaCl, 1 mM MgCl2, 1% Triton X-100, and complete protease inhibitor cocktail (Roche). To separate cell debris, the lysates were cleared by centrifugation at 4°C for 20 min at 16,000 g. Cell lysates were incubated with pre-blocked streptavidin beads for 120 min at 4°C followed by five washes with a buffer containing 20 mM Tris, pH 7.5, 150 mM NaCl, and 0.1% Triton X-100 (Hooikaas et al., 2019).

### Mass spectrometry analysis

Beads were re-suspended in 20 μL of Laemmli Sample buffer (Biorad) and supernatants were loaded on a 4-12% gradient Criterion XT Bis-Tris precast gel (Biorad). The gel was fixed with 40% methanol/10% acetic acid and then stained for 1 h using colloidal coomassie dye G-250 (Gel Code Blue Stain Reagent, Thermo Scientific). After in-gel digestion (Noordstra et al., 2016), all samples were analyzed on a Orbitrap Q-Exactive HF mass spectrometer (Thermo Scientific) coupled to an Agilent 1290 Infinity LC (Agilent Technologies). Peptides were loaded onto a trap column (Reprosil C18, 3 μm, 2 cm × 100 μm; Dr. Maisch) with solvent A (0.1% formic acid in water) at a maximum pressure of 800 bar and chromatographically separated over the analytical column (Zorbax SB-C18, 1.8 μm, 40 cm × 50 μm; Agilent) using 90 min linear gradient from 7-30% solvent B (0.1% formic acid in acetonitrile) at a flow rate of 150 nL/min. The mass spectrometer was used in a data-dependent mode, which automatically switched between MS and MS/MS. After a survey scan from 350-1500 m/z the 12 most abundant peptides were subjected to HCD fragmentation. For data analysis, raw files were processed using Proteome Discoverer 1.4 (Thermo Scientific). Database searches were performed using Mascot as search engine on the Human Uniprot database. Carbamidomethylation of cysteines was set as a fixed modification and oxidation of methionine was set as a variable modification. Trypsin was set as cleavage specificity, allowing a maximum of 2 missed cleavages. Data filtering was performed using percolator, resulting in 1% false discovery rate (FDR). Additional filters were search engine rank 1 and mascot ion score >20. To retrieve significant putative binding partners of Cfap53 additional analyses on the raw MS data were performed using Contaminant Repository for Affinity Purification (the CRAPome) implementing fold change scoring (FC) scoring and probalistic scoring using SAINT. The FC is calculated based on the ratio of average normalized spectral counts in bait purifications to negative controls. SAINT uses statistical modeling which provides a probability of true interaction (Choi et al., 2011; Mellacheruvu et al., 2013). The FC scoring is subdivided into a FCA (primary score based on BirA negative control pull down) and FCB (secondary score based on existing GFP negative control). The FCA and FCB cut-off were set to seven and three. The SAINT probability cut-off was set to 0.95.

### Drug treatments

To destabilize or destabilize the microtubule network 4 μg/ml nocodazole (1 mg/ml stock dissolved in DMSO) and 10 μM Taxol (1mM stock dissolved in DMSO) were used. To inhibit actin 25 μM Latrunculin A (5 mM stock dissolved in DMSO) was used. Embryos from the same fish were collected directly after fertilization and treated with the drug or DMSO control for 45 minutes, corresponding with the duration of the first cell division. Embryos were subsequently washed in E3 medium prior to fixation in microtubule fix as below and stained for γ-tubulin.

### mRNA Transcription and Injection

The PCS2-GFP-Cfap53 construct was linearized with *NotI* and transcribed and capped using the mMessage (SP6 kit Ambion, catalog # AM1340, Singapore). mRNA was purified using Lithium Chloride precipitation and re-suspension in nuclease free water. mRNA was injected into the yolk of 1-cell stage embryos in a volume of 1 nl at a concentration of 100 ng/μl.

### Fixation and Antibody labeling

For immunolabeling, HeLa cells were fixed in −20°C methanol for 10 min. Cells were then permeabilized with 0.15% Triton X-100 in PBS for 2 min; subsequent wash steps were performed in PBS supplemented with 0.05% Tween-20. Epitope blocking and antibody labeling steps were performed in PBS supplemented with 0.05% Tween-20 and 1% BSA. Before mounting in Vectashield mounting medium (Vector Laboratories), slides were washed with 70% and 100% ethanol and air-dried (Hooikaas et al., 2019).

Zebrafish embryos were collected at the right stages during development and fixed for two hours at room temperature in microtubule fix (3.7% formaldehyde, 0.25% glutaraldehyde, 5 mM EGTA, 0.3% Triton X-100) at the indicated stages. After fixation, embryos were washed 3 times with PBS-triton-X-100 (0.3%, PBST) and manually dechorionated. Embryos were dehydrated in methanol over night at −20°C and rehydrated in a methanol:PBS series. Before labeling embryos were treated with 0.5 mg/ml sodium borohydrate for 30 minutes at room temperature to inactivate remaining glutaraldehyde. Subsequently, embryos were washed in PBST and yolks were manually separated from the cells using forceps. Embryos were blocked in 0.3% PBST, 2 % BSA and 10% sheep serum for 1 hour at room temperature. Commercial antibodies used were as followed: anti-PCM1 (Sigma, rabbit polyclonal, 1:200), γ-tubulin (Sigma, mouse monoclonal GTU-88, 1:500), anti-centrin (Millipore, monoclonal mouse IgG2aκ 1:200) and anti-GFP (Aves, chicken, 1:500). All primary antibodies were incubated overnight at 4°C. Primary antibodies were detected using Alexa fluor 488 (Invitrogen, goat anti mouse), Alexa fluor 488 (Invitrogen, goat anti chicken, IgG1) and Alexa fluor 555 (Invitrogen, goat anti mouse, IgG1). Nuclei were shown by DAPI (4’,6-diamidino-2-phenylindole) staining.

### Imaging

Fixed embryos were embedded in 0.25% agarose to be mounted on a glass bottom dish. Subsequently embryos were imaged using a Leica SP5 Multi Photon setup using a 25x water and 63x glycerol objective followed by a Z-stack maximum projection (step size 1 μm).

For generation of time lapse videos, live embryos were embedded in 0.25% agarose to be mounted on a glass bottom dish. Subsequently, embryos were imaged live using a temperature controlled AF7000 Widefield Fluorescence Microscope setup using a 10x dry objective maintaining the temperature at a constant 28°C. The interval between images was set to 2 minutes. Computation of colocalization was performed using the 3D colocalization tool in Bitplane Imaris software (V9.3.1).

### Sperm motility evaluation by Computer-Assisted Sperm Analysis (CASA)

Sperm cells were extracted from *cfap53 −/−* and wildtype males as previously described (Dooley et al., 2013). Directly after collection, sperm cells were transferred into 50ul of buffered sperm-motility inhibiting solution (BSMIS) on ice. Motility was assessed on the computer-assisted semen analysis (CASA) system Sperm Vision® (version 3.5, Minitüb, Tiefenbach, Germany). The system was operated at room temperature with an automated stage and four-chamber slides of 20 μm depth (Leja, Nieuw Vennep, the Netherlands). After sperm activation in tap water, a 3.0 μL aliquot was immediately transferred to each chamber. Motility assessments started approximately twelve seconds after motility activation. Spermatozoa were examined at 200 x magnification by a camera with a resolution of 648 x 484 pixels (Pulnix TM-6760CL, JAI A/S, Glostrup, Denmark). The Sperm Vision® software analyzed 15 successive fields in the central part of the chamber at a frame rate of 60 Hz per field. Acquisition time for each field was 0.5 seconds. The surface area for sperm head detection was set at 4 to 80 μm². The setup ensured that in the majority of samples (11 out of 14) more than 1000 spermatozoa could be analyzed (mean: 1897 sperm). Semen samples were assessed for the percentage of motile (total motility), and progressively motile spermatozoa (progressive motility). The following motility descriptors were recorded for each motile spermatozoon: straight line velocity (VSL), curved line velocity (VCL), average path velocity (VAP), linearity (LIN = VSL/VCL), straightness (STR = VSL/VAP), wobble (WOB = VAP/VCL), average amplitude of lateral head-displacement (ALH), and beat cross frequency (BCF). A spermatozoon was considered to be motile when it met one of the following three definitions; 1) average head orientation change (AOC) higher than 7° and BCF greater than 25 Hz, DSL greater than 3.5 μm, VSL greater than 8 μm/s and DSL greater than 15 μm, VAP greater than 15 μm/s. Significance between wild type and mutant spermatozoa was calculated using a student t-test (Table S1).

### Primers

#### *cfap53* genotyping

FW_*cfap53* 5’-TGTAAGGAGAAGGAAGCAGGA

RV_*cfap53* 3’-TCATCAATGCCCATCTGGTA

#### *cfap53* genotyping in GFP-cfap53 transgenic background

FW_*cfap53* GGTGCTGGAGTTCACCAAAG

RV_*cfap53* intron 3’-ccgttcagacCTTTCCCTCT

## Acknowledgements

We would like to thank Wouter Makkinje for the help with generating the GFP-Cfap53 transgenic line.

## Author contributions

S.W., F.T. and J.B. conceived the research. F.T. generated the zebrafish transgenic line. S.W., B.V., M.A., R.S., H.H. performed the experiments. S.W., B.V., R.S., H.H., analyzed the data together with J.B., F.T.. S.W and J.B. wrote, and F.T., B.V. A.A., H.H., B.A.J.R. revised the manuscript.

## Supplementary figures

**Figure S1.**
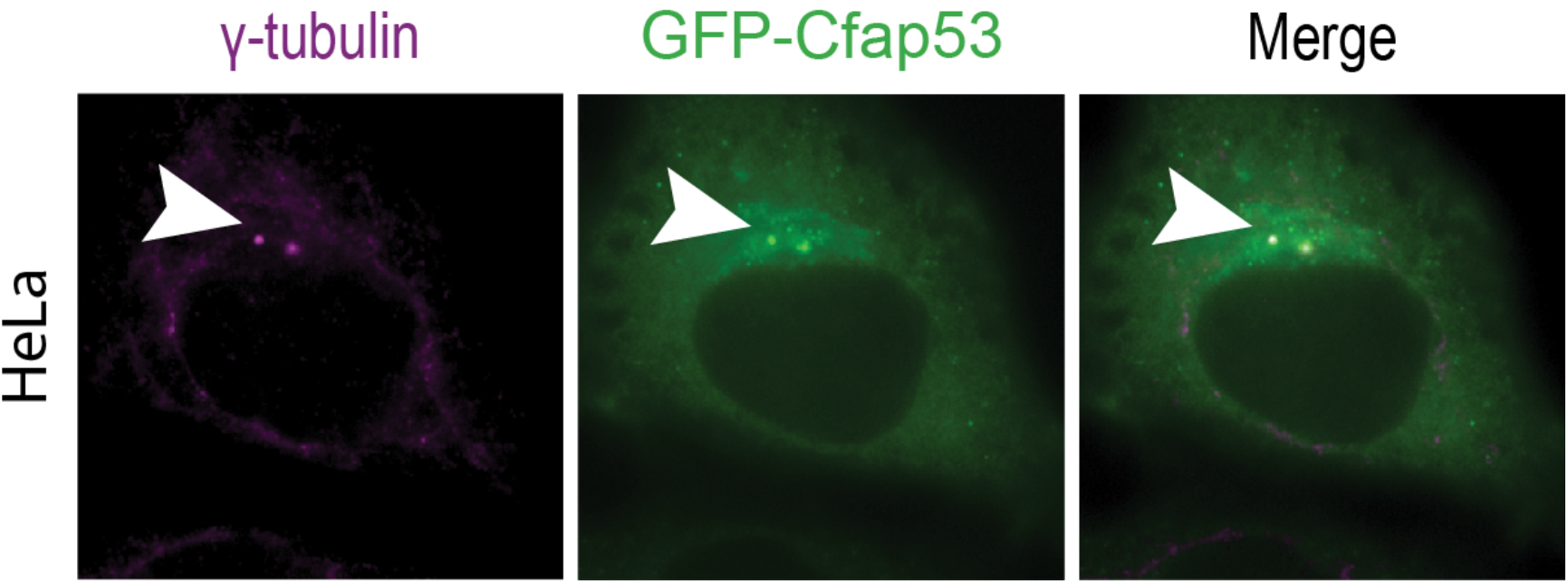
GFP-Cfap53 localization is centrosomal in HeLa transfected cells. GFP-Cfap53 has a centrosomal localization in transfected HeLa cells co-localizing with γ-tubulin indicated with arrowheads.

**Figure S2.**
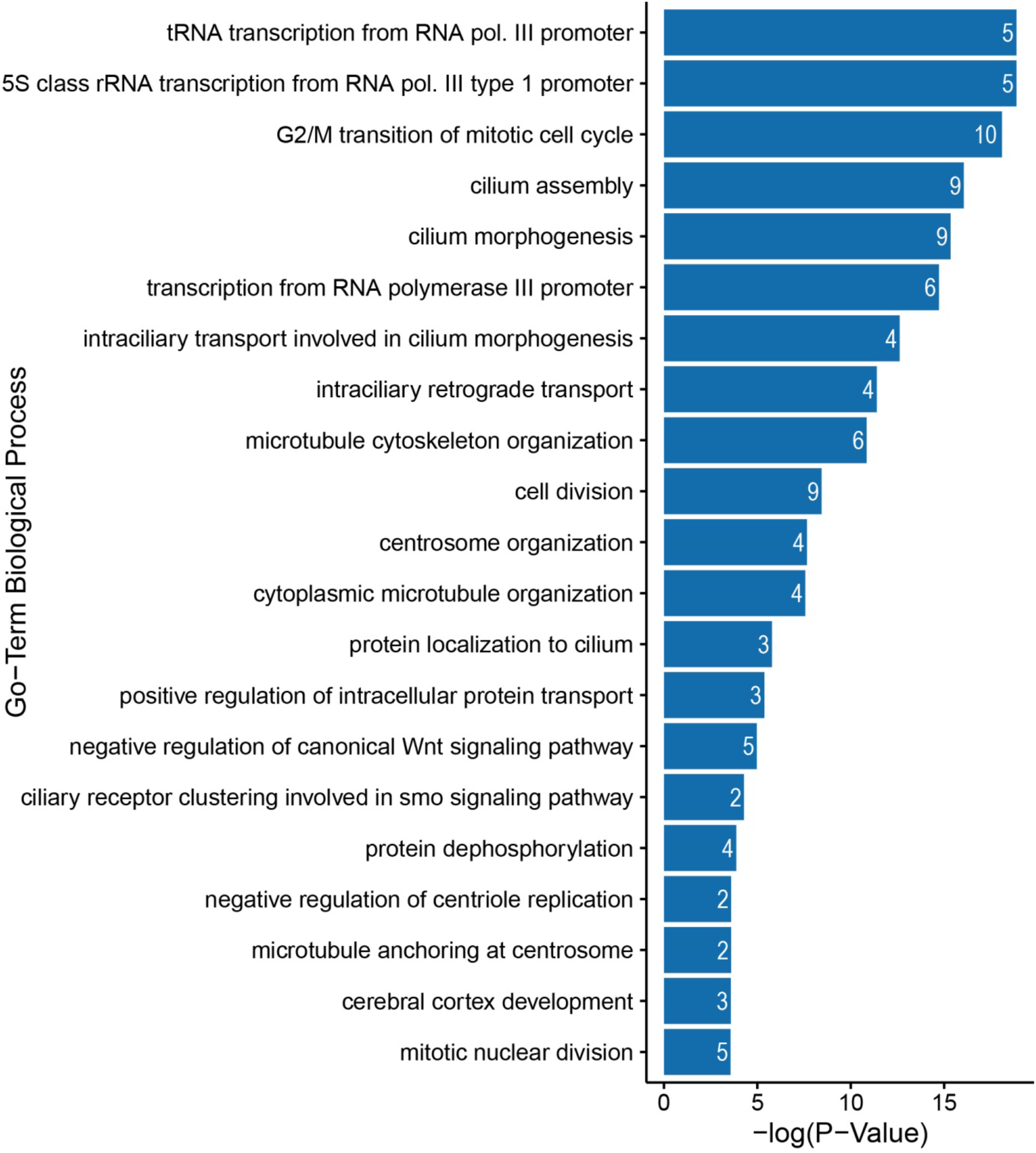
Functional GO analysis reveals a high overrepresentation of centrosomal, microtubule and cell cycle related biological processes. High confident hits from the MS data set were selected and analyzed using DAVID thereby indicating highly enriched GO-Term biological processes in our MS data set(Huang et al., 2009; Sherman and Lempicki, 2009). Cutoff p-value < 0.03.

**Figure S3.**
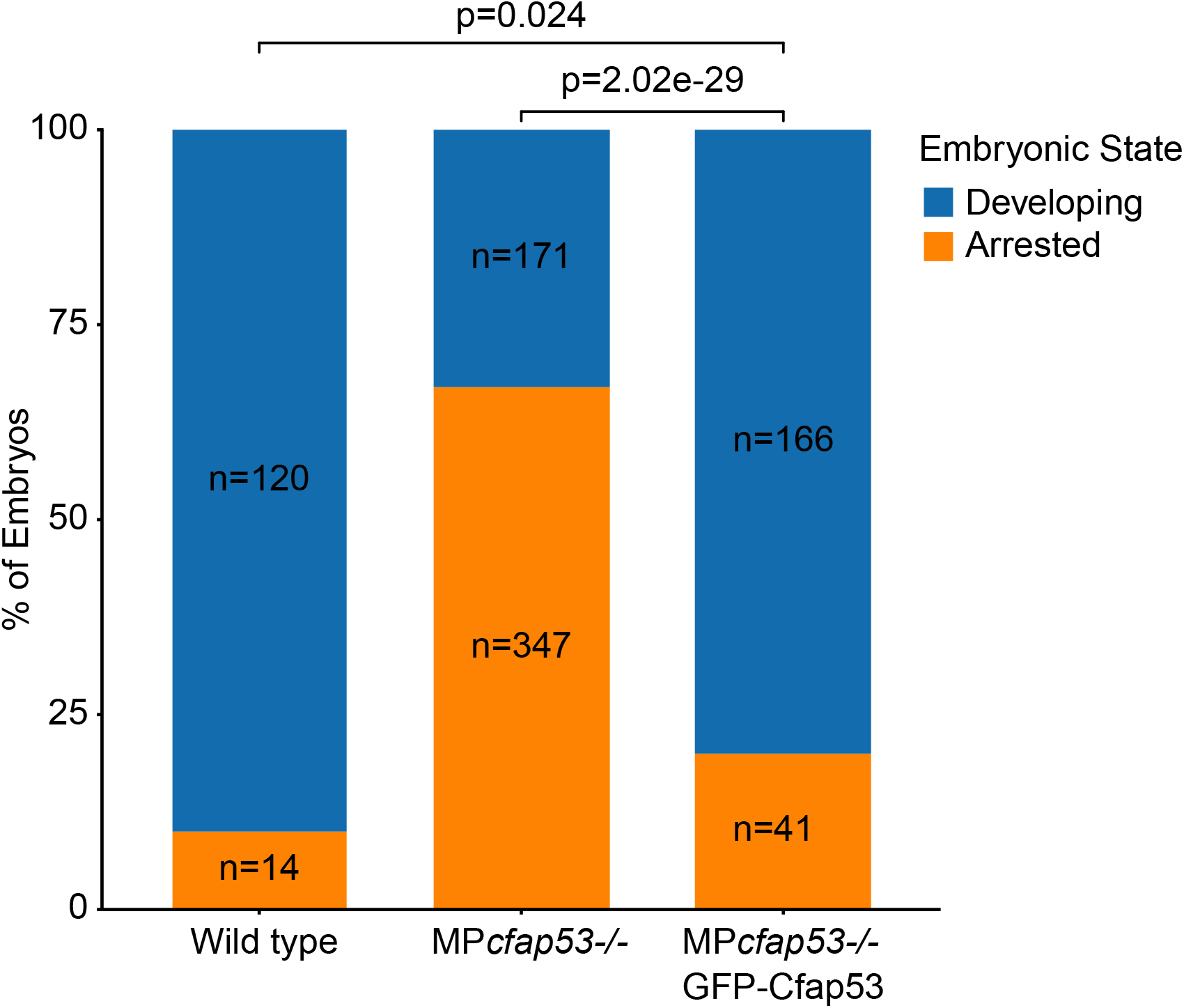
GFP-Cfap53 is a fully functional fusion protein. Bar plot indicating the number of developing and arrested embryos in clutches of wild type, MP*cfap53* −/− or Tg(*ubi:GFP-Cfap53*)/MP*cfap53*−/− embryos analyzed at 45 mpf. Chi-squared test was used to test for significance (p-value <= 0.05).

**Figure S4.**
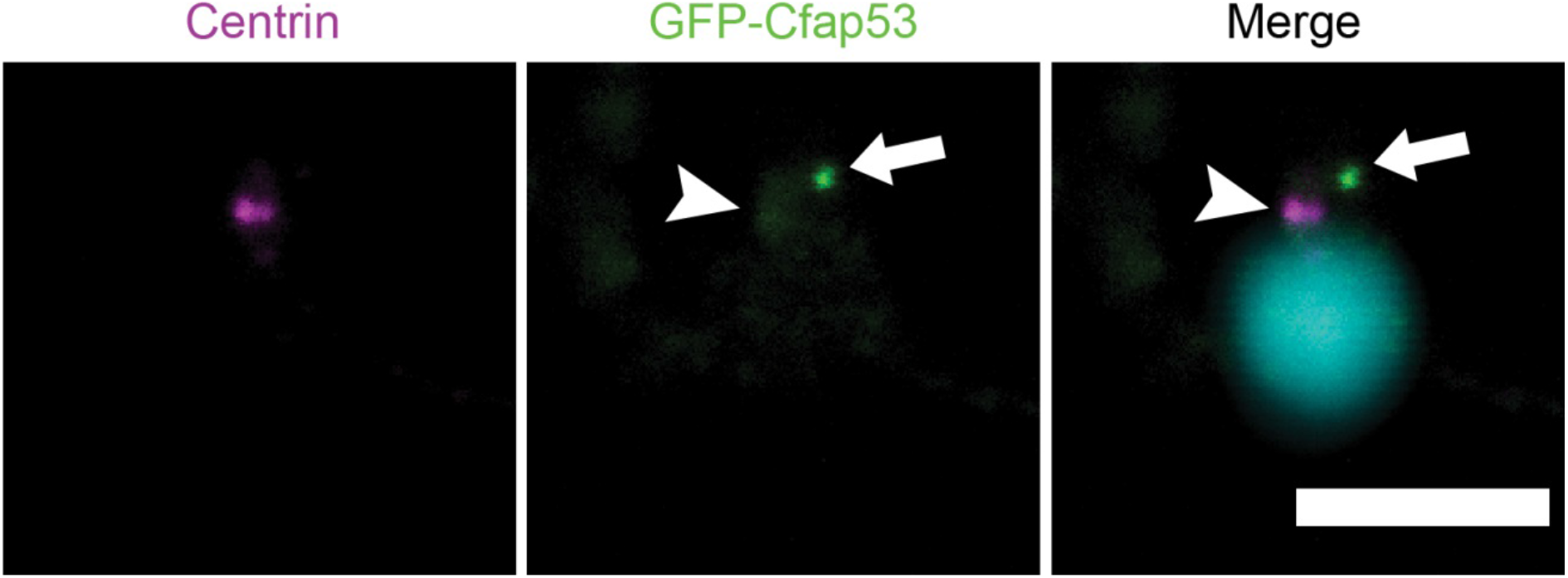
GFP-Cfap53 localization in sperm. Centrin and GFP immunolabeling in fixed zebrafish sperm cells. In the merge DAPI is shown in cyan. Arrowhead indicates mild GFP-Cfap53 colocalization with Centrin. Arrow indicates strong localization of GFP-Cfap53 in an uncharacterized structure in the sperm cell. Scalebar indicates 5 microns.

**Table S1.** Computer-assisted sperm analysis (CASA) of wild type and Cfap53 −/− sperm cells.

|  | **Fish ID** | **Genotype** | **Total number of sperm cells** | **% Motile sperm cells** |
| --- | --- | --- | --- | --- |
|  | 2 | mu | 2606 | 76.5 |
|  | 4 | mu | 1027 | 84.5 |
|  | 6 | mu | 414 | 83.1 |
|  | 8 | mu | 990 | 84.9 |
|  | 10 | mu | 2432 | 78.9 |
|  | 12 | mu | 2043 | 86.0 |
|  | 14 | mu | 1450 | 85.5 |
| **Mean** |  |  | 1566 | 83.0 |
| **SD** |  |  | 819 | 4.0 |
|  | 1 | wt | 1432 | 58.2 |
|  | 3 | wt | 381 | 61.9 |
|  | 5 | wt | 1052 | 87.1 |
|  | 7 | wt | 2492 | 88.4 |
|  | 9 | wt | 4981 | 83.5 |
|  | 11 | wt | 1689 | 84.0 |
|  | 13 | wt | 3565 | 87.8 |
| **Mean** |  |  | 2227 | 79 |
| **SD** |  |  | 1589 | 13 |
| **T-test (p-value)** |  |  | 0.347 | 0.437 |

**Table S2.** High confidence hits from GFP-CFAP53 and CFAP53-GFP AF-MS analysis.

| Accession | gene name | Score BirA | Score GFP-Cfap53 | Score Cfap53-GFP | FCA score | FCA>=7 | FCB score | FCB>=3 | SAINT probability | SAINT>=0.95 |
| --- | --- | --- | --- | --- | --- | --- | --- | --- | --- | --- |
| Q9Y5B8 | NME7 |  | 9712.88 | 9373.19 | 195.86 | TRUE | 105 | TRUE | 1 | TRUE |
| Q15154 | PCM1 |  | 4291.57 | 6558.99 | 145.35 | TRUE | 22.58 | TRUE | 1 | TRUE |
| O15078 | CEP290 |  | 3566.09 | 4333.98 | 136.39 | TRUE | 82.03 | TRUE | 1 | TRUE |
| Q12789 | GTF3C1 |  | 1472.83 | 5518.23 | 112.92 | TRUE | 19.37 | TRUE | 1 | TRUE |
| Q96M91 | CFAP53 | 232.85 | 73292.73 | 52087.98 | 77.77 | TRUE | 186.39 | TRUE | 1 | TRUE |
| Q14999 | CUL7 |  | 3572.68 | 2817.23 | 75.71 | TRUE | 36.12 | TRUE | 1 | TRUE |
| Q6P5Z2 | PKN3 |  | 4494.97 | 1531.99 | 74.23 | TRUE | 39.55 | TRUE | 1 | TRUE |
| Q8N4C6 | NIN |  | 2964.33 | 1122.01 | 66.11 | TRUE | 20.99 | TRUE | 1 | TRUE |
| Q9Y2I6 | NINL |  | 1668.40 | 1813.86 | 62.64 | TRUE | 37.82 | TRUE | 1 | TRUE |
| O94972 | TRIM37 |  | 1930.66 | 915.02 | 51.14 | TRUE | 30.32 | TRUE | 1 | TRUE |
| Q2M1P5 | KIF7 |  | 1343.00 | 1718.19 | 49.87 | TRUE | 30.27 | TRUE | 1 | TRUE |
| Q9UG01 | IFT172 |  | 1476.88 | 1245.93 | 47.08 | TRUE | 20.13 | TRUE | 1 | TRUE |
| A1XBS5 | FAM92A1 |  | 2409.88 | 2791.52 | 44.58 | TRUE | 27.1 | TRUE | 1 | TRUE |
| Q92621 | NUP205 | 34.08 | 2351.65 | 3213.88 | 44.07 | TRUE | 7.71 | TRUE | 1 | TRUE |
| Q9Y5Q9 | GTF3C3 |  | 1317.28 | 2326.24 | 43 | TRUE | 8.85 | TRUE | 1 | TRUE |
| Q9Y2K1 | ZBTB1 |  | 1866.79 | 1530.88 | 40.22 | TRUE | 24.46 | TRUE | 1 | TRUE |
| Q9Y5Q8 | GTF3C5 |  | 666.78 | 1521.12 | 36.45 | TRUE | 9.07 | TRUE | 1 | TRUE |
| Q9UKN8 | GTF3C4 |  | 527.51 | 2123.97 | 33.64 | TRUE | 8.32 | TRUE | 1 | TRUE |
| Q07864 | POLE |  | 428.18 | 1328.73 | 29.92 | TRUE | 10.67 | TRUE | 1 | TRUE |
| Q96RY7 | IFT140 |  | 888.15 | 658.20 | 29.03 | TRUE | 17.21 | TRUE | 1 | TRUE |
| Q5TB80 | CEP162 |  | 1131.49 | 472.55 | 28.72 | TRUE | 16.74 | TRUE | 1 | TRUE |
| Q6ZU80 | CEP128 |  | 622.24 | 506.35 | 28.4 | TRUE | 17.34 | TRUE | 1 | TRUE |
| A6NKD9 | CCDC85C |  | 1361.54 | 787.12 | 28.1 | TRUE | 16.68 | TRUE | 1 | TRUE |
| Q96C92 | SDCCAG3 |  | 2080.95 | 1198.38 | 27.48 | TRUE | 13.04 | TRUE | 1 | TRUE |
| O60318 | MCM3AP |  | 732.66 | 521.26 | 27.46 | TRUE | 13.31 | TRUE | 1 | TRUE |
| Q9Y5P8 | PPP2R3B |  | 877.37 | 945.72 | 25.59 | TRUE | 15.72 | TRUE | 1 | TRUE |
| Q9H6D7 | HAUS4 |  | 1016.32 | 555.32 | 24.99 | TRUE | 6.91 | TRUE | 1 | TRUE |
| Q9NVR7 | TBCCD1 |  | 913.02 | 993.49 | 24.97 | TRUE | 15.34 | TRUE | 1 | TRUE |
| Q13395 | TARBP1 |  | 1281.56 | 480.86 | 24.37 | TRUE | 10.25 | TRUE | 1 | TRUE |
| O95714 | HERC2 |  | 198.09 | 748.55 | 23.08 | TRUE | 13.01 | TRUE | 1 | TRUE |
| Q15058 | KIF14 |  | 569.83 | 417.42 | 22.8 | TRUE | 5.48 | TRUE | 1 | TRUE |
| Q9Y3M2 | CBY1 |  | 1311.47 | 1305.70 | 22.79 | TRUE | 14.04 | TRUE | 1 | TRUE |
| Q8TCG1 | KIAA1524 |  | 932.99 | 516.93 | 20.93 | TRUE | 12.58 | TRUE | 1 | TRUE |
| Q96JN8 | NEURL4 |  | 613.28 | 817.72 | 20.92 | TRUE | 12.88 | TRUE | 1 | TRUE |
| Q8WUA4 | GTF3C2 |  | 182.82 | 1341.30 | 20.89 | TRUE | 7.15 | TRUE | 1 | TRUE |
| O60232 | SSSCA1 |  | 2125.04 | 293.58 | 20.64 | TRUE | 7.06 | TRUE | 1 | TRUE |
| Q8NB25 | FAM184A |  | 728.91 | 408.52 | 20.63 | TRUE | 12.13 | TRUE | 1 | TRUE |
| Q8TD10 | MIPOL1 |  | 1092.86 | 565.77 | 20.31 | TRUE | 12.36 | TRUE | 1 | TRUE |
| Q5SW79 | CEP170 |  | 421.99 | 711.24 | 20.29 | TRUE | 3.67 | TRUE | 1 | TRUE |
| Q9UL15 | BAG5 |  | 1244.28 | 518.88 | 20.01 | TRUE | 9.23 | TRUE | 1 | TRUE |
| Q53T94 | TAF1B |  | 625.19 | 1085.98 | 19.67 | TRUE | 12.09 | TRUE | 1 | TRUE |
| Q9BV73 | CEP250 |  | 568.26 | 385.68 | 19.37 | TRUE | 10.49 | TRUE | 1 | TRUE |
| Q9UPN4 | CEP131 |  | 494.31 | 455.41 | 19.36 | TRUE | 6.99 | TRUE | 1 | TRUE |
| O14802 | POLR3A |  | 320.47 | 364.56 | 17.8 | TRUE | 6.13 | TRUE | 1 | TRUE |
| Q86VW0 | SESTD1 |  | 766.52 | 500.12 | 17.5 | TRUE | 10.87 | TRUE | 1 | TRUE |
| P42695 | NCAPD3 |  | 230.42 | 664.25 | 17.17 | TRUE | 7.04 | TRUE | 1 | TRUE |
| Q7Z4Q2 | HEATR3 |  | 603.73 | 494.12 | 16.56 | TRUE | 9.48 | TRUE | 1 | TRUE |
| Q13136 | PPFIA1 |  | 564.86 | 442.51 | 16.26 | TRUE | 7.98 | TRUE | 1 | TRUE |
| Q7Z7A1 | CNTRL |  | 591.36 | 194.10 | 15.96 | TRUE | 8.23 | TRUE | 1 | TRUE |
| Q96CS2 | HAUS1 |  | 892.15 | 247.33 | 15.95 | TRUE | 8.72 | TRUE | 1 | TRUE |
| Q12923 | PTPN13 |  | 298.28 | 606.29 | 15.93 | TRUE | 9.6 | TRUE | 1 | TRUE |
| Q15834 | CCDC85B |  | 523.45 | 259.31 | 15.33 | TRUE | 9.31 | TRUE | 1 | TRUE |
| O94964 | SOGA1 |  | 127.53 | 597.99 | 14.99 | TRUE | 6.44 | TRUE | 1 | TRUE |
| Q92574 | TSC1 |  | 829.10 | 229.59 | 14.71 | TRUE | 8.04 | TRUE | 1 | TRUE |
| Q8N3Y1 | FBXW8 |  | 608.23 | 810.83 | 14.69 | TRUE | 9.08 | TRUE | 1 | TRUE |
| Q709F0 | ACAD11 |  | 520.08 | 599.16 | 14.69 | TRUE | 9.08 | TRUE | 1 | TRUE |
| Q96SN8 | CDK5RAP2 |  | 297.33 | 346.72 | 14.39 | TRUE | 8.04 | TRUE | 1 | TRUE |
| Q9NQC7 | CYLD |  | 512.00 | 183.38 | 13.77 | TRUE | 8.59 | TRUE | 1 | TRUE |
| Q9UL42 | PNMA2 |  | 579.18 | 357.88 | 13.77 | TRUE | 8.3 | TRUE | 1 | TRUE |
| Q8NEZ3 | WDR19 |  | 356.13 | 233.78 | 13.45 | TRUE | 8.42 | TRUE | 1 | TRUE |
| P42345 | MTOR |  | 169.75 | 178.28 | 13.14 | TRUE | 4.12 | TRUE | 1 | TRUE |
| Q86SQ0 | PHLDB2 |  | 549.97 | 177.69 | 12.53 | TRUE | 7.06 | TRUE | 1 | TRUE |
| Q9H1A4 | ANAPC1 |  | 235.49 | 245.08 | 11.89 | TRUE | 7.5 | TRUE | 1 | TRUE |
| Q8IWT3 | CUL9 |  | 650.78 | 583.43 | 11.89 | TRUE | 4.92 | TRUE | 1 | TRUE |
| P23258 | TUBG1 |  | 509.59 | 288.73 | 11.59 | TRUE | 4.89 | TRUE | 1 | TRUE |
| Q7Z494 | NPHP3 |  | 594.04 | 26.64 | 11.29 | TRUE | 4.68 | TRUE | 1 | TRUE |
| Q8TED0 | UTP15 |  | 131.05 | 455.84 | 10.95 | TRUE | 5.61 | TRUE | 1 | TRUE |
| Q7Z4L5 | TTC21B |  | 254.32 | 147.79 | 10.66 | TRUE | 6.46 | TRUE | 1 | TRUE |
| P07199 | CENPB |  | 519.75 | 331.42 | 10.35 | TRUE | 3.32 | TRUE | 1 | TRUE |
| Q06190 | PPP2R3A |  | 290.99 | 276.44 | 10.34 | TRUE | 6.59 | TRUE | 1 | TRUE |
| Q9UPU5 | USP24 |  | 376.51 | 133.43 | 10.03 | TRUE | 4.36 | TRUE | 1 | TRUE |
| Q96GE4 | CEP95 |  | 316.56 | 288.25 | 10.02 | TRUE | 6.38 | TRUE | 1 | TRUE |
| Q9HD67 | MYO10 |  | 201.65 | 201.65 | 9.71 | TRUE | 6.21 | TRUE | 1 | TRUE |
| Q9C099 | LRRCC1 |  | 215.78 | 89.83 | 9.42 | TRUE | 4.89 | TRUE | 1 | TRUE |
| Q9HBG6 | IFT122 |  | 264.23 | 136.12 | 8.47 | TRUE | 5.36 | TRUE | 1 | TRUE |
| Q9BWT7 | CARD10 |  | 153.06 | 119.83 | 8.16 | TRUE | 5.26 | TRUE | 1 | TRUE |
| Q15029 | EFTUD2 |  | 181.21 | 282.05 | 8.15 | TRUE | 4.23 | TRUE | 1 | TRUE |
| Q8WV44 | TRIM41 |  | 64.95 | 368.58 | 8.15 | TRUE | 3.31 | TRUE | 1 | TRUE |
| A1A4S6 | ARHGAP10 |  | 382.43 | 32.38 | 7.86 | TRUE | 3.84 | TRUE | 1 | TRUE |
| P14373 | TRIM27 |  | 283.78 | 223.57 | 7.85 | TRUE | 3.36 | TRUE | 1 | TRUE |
| Q9Y4C2 | TCAF1 |  | 240.29 | 169.07 | 7.54 | TRUE | 4.91 | TRUE | 1 | TRUE |
| Q5JTC6 | AMER1 |  | 283.72 | 214.16 | 7.54 | TRUE | 3.67 | TRUE | 1 | TRUE |
| Q8NCN5 | PDPR |  | 308.17 | 126.01 | 7.23 | TRUE | 4.73 | TRUE | 1 | TRUE |
| Q06481 | APLP2 |  | 275.02 | 113.53 | 7.23 | TRUE | 4.67 | TRUE | 1 | TRUE |
| O00743 | PPP6C |  | 219.78 | 106.47 | 7.23 | TRUE | 4.59 | TRUE | 1 | TRUE |
| Q9GZT3 | SLIRP |  | 270.85 | 142.26 | 7.23 | TRUE | 4.35 | TRUE | 1 | TRUE |
| Q96RT7 | TUBGCP6 |  | 237.52 | 201.74 | 7.23 | TRUE | 3.97 | TRUE | 1 | TRUE |
| P51530 | DNA2 |  | 153.36 | 178.44 | 7.22 | TRUE | 4.71 | TRUE | 1 | TRUE |

**Table S3.**
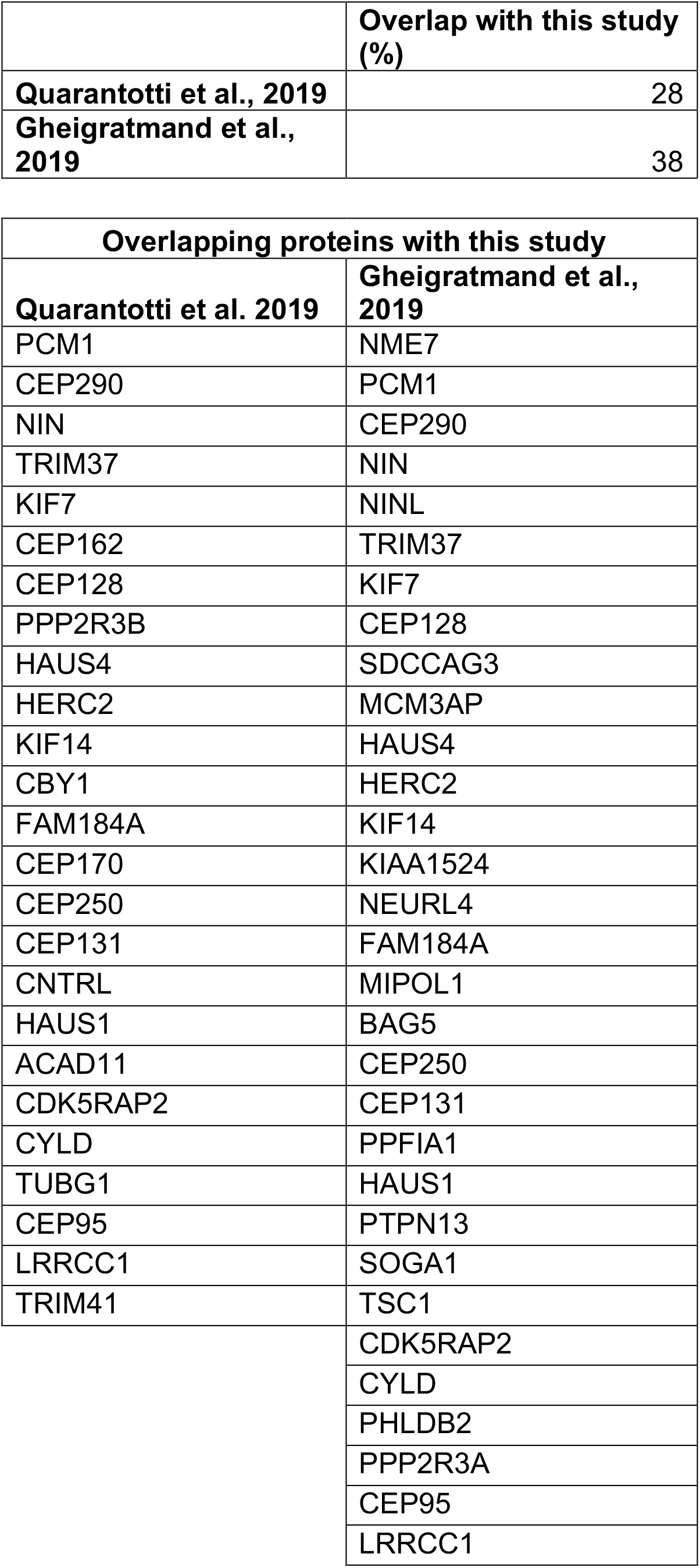
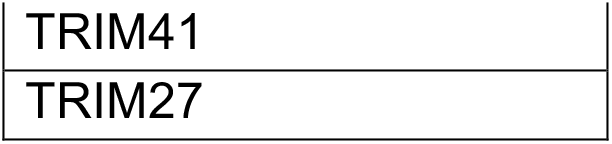
Overlap of identified proteins in this study compared to two centriolar satellite proteomic studies.

|  | **Overlap with this study (%)** |
| --- | --- |
| **Quarantotti et al., 2019** | 28 |
| **Gheigratmand et al., 2019** | 38 |

| **Overlapping proteins with this study** |  |
| --- | --- |
| **Quarantotti et al. 2019** | **Gheigratmand et al., 2019** |
| PCM1 | NME7 |
| CEP290 | PCM1 |
| NIN | CEP290 |
| TRIM37 | NIN |
| KIF7 | NINL |
| CEP162 | TRIM37 |
| CEP128 | KIF7 |
| PPP2R3B | CEP128 |
| HAUS4 | SDCCAG3 |
| HERC2 | MCM3AP |
| KIF14 | HAUS4 |
| CBY1 | HERC2 |
| FAM184A | KIF14 |
| CEP170 | KIAA1524 |
| CEP250 | NEURL4 |
| CEP131 | FAM184A |
| CNTRL | MIPOL1 |
| HAUS1 | BAG5 |
| ACAD11 | CEP250 |
| CDK5RAP2 | CEP131 |
| CYLD | PPFIA1 |
| TUBG1 | HAUS1 |
| CEP95 | PTPN13 |
| LRRCC1 | SOGA1 |
| TRIM41 | TSC1 |
|  | CDK5RAP2 |
|  | CYLD |
|  | PHLDB2 |
|  | PPP2R3A |
|  | CEP95 |
|  | LRRCC1 |

|  |
| --- |
| TRIM41 |
| TRIM27 |

## References

Abrams, E. W., Fuentes, R., Marlow, F. L., Kobayashi, M., Zhang, H., Lu, S., Kapp, L., Joseph, S. R., Kugath, A. and Gupta, T. (2020). Molecular genetics of maternally-controlled cell divisions. PLoS Genetics 16, e1008652.

Aleström, P., D’Angelo, L., Midtlyng, P. J., Schorderet, D. F., Schulte-Merker, S., Sohm, F. and Warner, S. (2020). Zebrafish: Housing and husbandry recommendations. Laboratory Animals 54, 213–224.

Bakkers, J., Verhoeven, M. C. and Abdelilah-Seyfried, S. (2009). Shaping the zebrafish heart: From left–right axis specification to epithelial tissue morphogenesis. Developmental biology 330, 213–220.

Balczon, R., Bao, L. and Zimmer, W. E. (1994). PCM-1, A 228-kD centrosome autoantigen with a distinct cell cycle distribution. The Journal of cell biology 124, 783–793.

Choi, H., Larsen, B., Lin, Z.-Y., Breitkreutz, A., Mellacheruvu, D., Fermin, D., Qin, Z. S., Tyers, M., Gingras, A.-C. and Nesvizhskii, A. I. (2011). SAINT: probabilistic scoring of affinity purification–mass spectrometry data. Nature methods 8, 70–73.

Choi, Y.-K., Liu, P., Sze, S. K., Dai, C. and Qi, R. Z. (2010). CDK5RAP2 stimulates microtubule nucleation by the γ-tubulin ring complex. The Journal of cell biology 191, 1089–1095.

Conkar, D., Bayraktar, H. and Firat-Karalar, E. N. (2019). Centrosomal and ciliary targeting of CCDC66 requires cooperative action of centriolar satellites, microtubules and molecular motors. Scientific Reports 9, 14250–14250.

Dammermann, A. and Merdes, A. (2002). Assembly of centrosomal proteins and microtubule organization depends on PCM-1. The Journal of cell biology 159, 255–266.

Dekens, M. P. S., Pelegri, F. J., Maischein, H.-M. and Nüsslein-Volhard, C. (2003). The maternal-effect gene futile cycle is essential for pronuclear congression and mitotic spindle assembly in the zebrafish zygote. Development 130, 3907–3916.

Dooley, C. M., Scahill, C., Fényes, F., Kettleborough, R. N., Stemple, D. L. and Busch-Nentwich, E. M. (2013). Multi-allelic phenotyping–a systematic approach for the simultaneous analysis of multiple induced mutations. Methods 62, 197–206.

Doxsey, S. (2001). Re-evaluating centrosome function. Nature reviews molecular cell biology 2, 688–698.

Félix, M. A., Antony, C., Wright, M. and Maro, B. (1994). Centrosome assembly in vitro: role of gamma-tubulin recruitment in Xenopus sperm aster formation. The Journal of cell biology 124, 19–31.

Firat-Karalar, E. N., Sante, J., Elliott, S. and Stearns, T. (2014). Proteomic analysis of mammalian sperm cells identifies new components of the centrosome. Journal of cell science 127, 4128–4133.

Gheiratmand, L., Coyaud, E., Gupta, G. D., Laurent, E. M., Hasegan, M., Prosser, S. L., Gonçalves, J., Raught, B. and Pelletier, L. (2019). Spatial and proteomic profiling reveals centrosome-independent features of centriolar satellites. The EMBO journal 38, e101109.

Gillingham, A. K. and Munro, S. (2000). The PACT domain, a conserved centrosomal targeting motif in the coiled-coil proteins AKAP450 and pericentrin. EMBO reports 1, 524–529.

Goshima, G., Mayer, M., Zhang, N., Stuurman, N. and Vale, R. D. (2008). Augmin: a protein complex required for centrosome-independent microtubule generation within the spindle. The Journal of cell biology 181, 421–429.

Gould, R. R. and Borisy, G. G. (1977). The pericentriolar material in Chinese hamster ovary cells nucleates microtubule formation. The Journal of cell biology 73, 601–615.

Gueth-Hallonet, C., Antony, C., Aghion, J., Santa-Maria, A., Lajoie-Mazenc, I., Wright, M. and Maro, B. (1993). gamma-Tubulin is present in acentriolar MTOCs during early mouse development. Journal of cell science 105, 157–166.

Hirokawa, N., Tanaka, Y., Okada, Y. and Takeda, S. (2006). Nodal flow and the generation of left-right asymmetry. Cell 125, 33–45.

Holy, J. and Schatten, G. (1997). Recruitment of maternal material during assembly of the zygote centrosome in fertilized sea urchin eggs. Cell and tissue research 289, 285–297.

Hooikaas, P. J., Martin, M., Mühlethaler, T., Kuijntjes, G.-J., Peeters, C. A., Katrukha, E. A., Ferrari, L., Stucchi, R., Verhagen, D. G. and Van Riel, W. E. (2019). MAP7 family proteins regulate kinesin-1 recruitment and activation. The Journal of cell biology 218, 1298–1318.

Hori, A. and Toda, T. (2017). Regulation of centriolar satellite integrity and its physiology. Cellular and Molecular Life Sciences 74, 213–229.

Huang, D. W., Sherman, B. T. and Lempicki, R. A. (2009). Bioinformatics enrichment tools: paths toward the comprehensive functional analysis of large gene lists. Nucleic acids research 37, 1–13.

Hutchins, J. R., Toyoda, Y., Hegemann, B., Poser, I., Hériché, J.-K., Sykora, M. M., Augsburg, M., Hudecz, O., Buschhorn, B. A. and Bulkescher, J. (2010). Systematic analysis of human protein complexes identifies chromosome segregation proteins. Science 328, 593–599.

Inoue, D., Stemmer, M., Thumberger, T., Ruppert, T., Bärenz, F., Wittbrodt, J. and Gruss, O. J. (2017). Expression of the novel maternal centrosome assembly factor Wdr8 is required for vertebrate embryonic mitoses. Nature communications 8, 14090–14090.

Jia, Y., Fong, K. W., Choi, Y. K., See, S. S. and Qi, R. Z. (2013). Dynamic Recruitment of CDK5RAP2 to Centrosomes Requires Its Association with Dynein. PLoS ONE 8, e68523.

Khodjakov, A. and Rieder, C. L. (1999). The sudden recruitment of gamma-tubulin to the centrosome at the onset of mitosis and its dynamic exchange throughout the cell cycle, do not require microtubules. The Journal of cell biology 146, 585–596.

Kim, J., Krishnaswami, S. R. and Gleeson, J. G. (2008). CEP290 interacts with the centriolar satellite component PCM-1 and is required for Rab8 localization to the primary cilium. Human molecular genetics 17, 3796–3805.

Kimmel, C. B., Ballard, W. W., Kimmel, S. R., Ullmann, B. and Schilling, T. F. (1995). Stages of embryonic development of the zebrafish. Developmental dynamics 203, 253–310.

Kollman, J. M., Merdes, A., Mourey, L. and Agard, D. A. (2011). Microtubule nucleation by γ-tubulin complexes. Nature reviews Molecular cell biology 12, 709–721.

Kubo, A., Sasaki, H., Yuba-Kubo, A., Tsukita, S. and Shiina, N. (1999). Centriolar satellites: Molecular characterization, ATP-dependent movement toward centrioles and possible involvement in ciliogenesis. The Journal of cell biology 147, 969–979.

Kubo, A. and Tsukita, S. (2003). Non-membranous granular organelle consisting of PCM-1: subcellular distribution and cell-cycle-dependent assembly/disassembly. Journal of cell science 116, 919–928.

Kwan, K. M., Fujimoto, E., Grabher, C., Mangum, B. D., Hardy, M. E., Campbell, D. S., Parant, J. M., Yost, H. J., Kanki, J. P. and Chien, C. B. (2007). The Tol2kit: a multisite gateway-based construction kit for Tol2 transposon transgenesis constructs. Developmental dynamics 236, 3088–3099.

Lawo, S., Bashkurov, M., Mullin, M., Ferreria, M. G., Kittler, R., Habermann, B., Tagliaferro, A., Poser, I., Hutchins, J. R. and Hegemann, B. (2009). HAUS, the 8-subunit human Augmin complex, regulates centrosome and spindle integrity. Current biology 19, 816–826.

Lindeman, R. E. and Pelegri, F. (2012). Localized products of futile cycle/ lrmp promote centrosome-nucleus attachment in the zebrafish zygote. Current biology 22, 843–851.

Liu, P., Choi, Y.-K. and Qi, R. Z. (2014). NME7 is a functional component of the γ-tubulin ring complex. Molecular biology of the cell 25, 2017–2025.

Mason, J. M. and Arndt, K. M. (2004). Coiled Coil Domains: Stability, Specificity, and Biological Implications. ChemBioChem 5, 170–176.

Mellacheruvu, D., Wright, Z., Couzens, A. L., Lambert, J. P., St-Denis, N. A., Li, T., Miteva, Y. V., Hauri, S., Sardiu, M. E., Low, T. Y., et al. (2013). The CRAPome: A contaminant repository for affinity purification-mass spectrometry data. Nature Methods 10, 730–736.

Moritz, M., Zheng, Y., Alberts, B. M. and Oegema, K. (1998). Recruitment of the γ-tubulin ring complex to Drosophila salt-stripped centrosome scaffolds. The journal of cell biology 142, 775–786.

Mosimann, C., Kaufman, C. K., Li, P., Pugach, E. K., Tamplin, O. J. and Zon, L. I. (2011). Ubiquitous transgene expression and Cre-based recombination driven by the ubiquitin promoter in zebrafish. Development 138, 169–177.

Murphy, S. M., Preble, A. M., Patel, U. K., O’Connell, K. L., Dias, D. P., Moritz, M., Agard, D., Stults, J. T. and Stearns, T. (2001). GCP5 and GCP6: two new members of the human γ-tubulin complex. Molecular biology of the cell 12, 3340–3352.

Narasimhan, V., Hjeij, R., Vij, S., Loges, N. T., Wallmeier, J., Koerner-Rettberg, C., Werner, C., Thamilselvam, S. K., Boey, A., Choksi, S. P., et al. (2015). Mutations in CCDC11, which encodes a coiled-coil containing ciliary protein, causes situs inversus due to dysmotility of monocilia in the left-right organizer. Human mutation 36, 307–318.

Noël, E. S., Momenah, T. S., Al-Dagriri, K., Al-Suwaid, A., Al-Shahrani, S., Jiang, H., Willekers, S., Oostveen, Y. Y., Chocron, S., Postma, A. V., et al. (2016). A zebrafish Loss-of-Function Model for Human CFAP53 Mutations Reveals its Specific Role in Laterality Organ Function. Human mutation 37, 194–200.

Noordstra, I., Liu, Q., Nijenhuis, W., Hua, S., Jiang, K., Baars, M., Remmelzwaal, S., Martin, M., Kapitein, L. C. and Akhmanova, A. (2016). Control of apico– basal epithelial polarity by the microtubule minus-end-binding protein CAMSAP3 and spectraplakin ACF7. Journal of cell science 129, 4278–4288.

Odabasi, E., Gul, S., Kavakli, I. H. and Firat-Karalar, E. N. (2019). Centriolar satellites are required for efficient ciliogenesis and ciliary content regulation. EMBO reports 20, e47723.

Pauls, S., Geldmacher-Voss, B. and Campos-Ortega, J. A. (2001). A zebrafish histone variant H2A. F/Z and a transgenic H2A. F/Z: GFP fusion protein for in vivo studies of embryonic development. Development genes and evolution 211, 603–610.

Pelegri, F. and Mullins, M. (2016). Genetic screens for mutations affecting adult traits and parental-effect genes. In Methods in Cell Biology, pp. 39–87: Elsevier.

Perles, Z., Cinnamon, Y., Ta-Shma, A., Shaag, A., Einbinder, T., Rein, A. J. J. T. and Elpeleg, O. (2012). A human laterality disorder associated with recessive CCDC11 mutation. Journal of medical genetics 49, 386–390.

Prosser, S. L. and Pelletier, L. (2020). Centriolar satellite biogenesis and function in vertebrate cells. Journal of cell science 133, jcs239566.

Quarantotti, V., Chen, J. X., Tischer, J., Gonzalez Tejedo, C., Papachristou, E. K., D’Santos, C. S., Kilmartin, J. V., Miller, M. L. and Gergely, F. (2019). Centriolar satellites are acentriolar assemblies of centrosomal proteins. The EMBO Journal 38, e101082.

Quintyne, N., Gill, S., Eckley, D., Crego, C., Compton, D. and Schroer, T. (1999). Dynactin is required for microtubule anchoring at centrosomes. The Journal of cell biology 147, 321–334.

Rathbun, L., Aljiboury, A. A., Bai, X., Manikas, J., Amack, J. D., Bembenek, J. N. and Hehnly, H. (2020). PLK1-and PLK4-mediated asymmetric mitotic centrosome size and positioning in the early zebrafish embryo. Current biology.

Reichmann, J., Nijmeijer, B., Hossain, M. J., Eguren, M., Schneider, I., Politi, A. Z., Roberti, M. J., Hufnagel, L., Hiiragi, T. and Ellenberg, J. (2018). Dual-spindle formation in zygotes keeps parental genomes apart in early mammalian embryos. Science 361, 189–193.

Rossi, A., Kontarakis, Z., Gerri, C., Nolte, H., Hölper, S., Krüger, M. and Stainier, D. Y. (2015). Genetic compensation induced by deleterious mutations but not gene knockdowns. Nature 524, 230–233.

Schatten, G. (1994). The centrosome and its mode of inheritance: The reduction of the centrosome during gametogenesis and its restoration during fertilization. Developmental biology 165, 299–335.

Schatten, H., Schatten, G., Mazia, D., Balczon, R. and Simerly, C. (1986). Behavior of centrosomes during fertilization and cell division in mouse oocytes and in sea urchin eggs. Proceedings of the National Academy of Sciences 83, 105–109.

Schatten, H., Thompson-Coffe, C., Coffe, G., Simerly, C. and Schatten, G. (1989). Centrosomes, centrioles, and posttranslationally modified α-tubulins during fertilization. In The molecular biology of fertilization, pp. 189–210.

Schnackenberg, B. J., Khodjakov, A., Rieder, C. L. and Palazzo, R. E. (1998). The disassembly and reassembly of functional centrosomes in vitro. Proceedings of the National Academy of Sciences 95, 9295–9300.

Sherman, B. T. and Lempicki, R. A. (2009). Systematic and integrative analysis of large gene lists using DAVID bioinformatics resources. Nature protocols 4, 44.

Silva, E., Betleja, E., John, E., Spear, P., Moresco, J. J., Zhang, S., Yates, J. R., Mitchell, B. J. and Mahjoub, M. R. (2016). Ccdc11 is a novel centriolar satellite protein essential for ciliogenesis and establishment of left–right asymmetry. Molecular biology of the cell 27, 48–63.

Staples, C. J., Myers, K. N., Beveridge, R. D., Patil, A. A., Lee, A. J., Swanton, C., Howell, M., Boulton, S. J. and Collis, S. J. (2012). The centriolar satellite protein Cep131 is important for genome stability. Journal of cell science 125, 4770–4779.

Stearns, T. and Kirschner, M. (1994). In vitro reconstitution of centrosome assembly and function: the central role of gamma-tubulin. Cell 76, 623–637.

Stowe, T. R., Wilkinson, C. J., Iqbal, A. and Stearns, T. (2012). The centriolar satellite proteins Cep72 and Cep290 interact and are required for recruitment of BBS proteins to the cilium. Molecular biology of the cell 23, 3322–3335.

Sutovsky, P. and Schatten, G. (1999). Paternal contributions to the mammalian zygote: fertilization after sperm-egg fusion. International review of cytology 195, 1–65.

Szklarczyk, D., Franceschini, A., Wyder, S., Forslund, K., Heller, D., Huerta-Cepas, J., Simonovic, M., Roth, A., Santos, A. and Tsafou, K. P. (2015). STRING v10: protein–protein interaction networks, integrated over the tree of life. Nucleic acids research 43, D447–D452.

Szollosi, D., Calarco, P. and Donahue, R. (1972). Absence of centrioles in the first and second meiotic spindles of mouse oocytes. Journal of cell science 11, 521–541.

Sztal, T. E., McKaige, E. A., Williams, C., Ruparelia, A. A. and Bryson-Richardson, R. J. (2018). Genetic compensation triggered by actin mutation prevents the muscle damage caused by loss of actin protein. PLoS genetics 14, e1007212.

Tran, L. D., Hino, H., Quach, H., Lim, S., Shindo, A., Mimori-Kiyosue, Y., Mione, M., Ueno, N., Winkler, C. and Hibi, M. (2012). Dynamic microtubules at the vegetal cortex predict the embryonic axis in zebrafish. Development 139, 3644–3652.

Woodruff, J. B. (2018). Assembly of mitotic structures through phase separation. Journal of molecular biology 430, 4762–4772.

Woodruff, J. B., Gomes, B. F., Widlund, P. O., Mahamid, J., Honigmann, A. and Hyman, A. A. (2017). The centrosome is a selective condensate that nucleates microtubules by concentrating tubulin. Cell 169, 1066–1077.

Woolley, D. M. and Fawcett, D. W. (1973). The degeneration and disappearance of the centrioles during the development of the rat spermatozoon. The Anatomical record 177, 289–301.

Yabe, T., Ge, X. and Pelegri, F. (2007). The zebrafish maternal-effect gene cellular atoll encodes the centriolar component sas-6 and defects in its paternal function promote whole genome duplication. Developmental biology 312, 44–60.

Young, A., Dictenberg, J. B., Purohit, A., Tuft, R. and Doxsey, S. J. (2000). Cytoplasmic dynein-mediated assembly of pericentrin and γ tubulin onto centrosomes. Molecular biology of the cell 11, 2047–2056.

Zhu, P., Ma, Z., Guo, L., Zhang, W., Zhang, Q., Zhao, T., Jiang, K., Peng, J. and Chen, J. (2017). Short body length phenotype is compensated by the upregulation of nidogen family members in a deleterious nid1a mutation of zebrafish. J. Genet. Genomics 44, 553–556.

Zimmerman, W. and Doxsey, S. J. (2000). Construction of centrosomes and spindle poles by molecular motor-driven assembly of protein particles. Traffic 1, 927–934.

